# Biotrophic interactions disentangled: *In situ* localisation of mRNAs to decipher plant and algal pathogen – host interactions at the single cell level

**DOI:** 10.1101/378794

**Authors:** Julia Badstöber, Claire M. M. Gachon, Jutta Ludwig-Müller, Adolf M. Sandbichler, Sigrid Neuhauser

## Abstract

Plant-pathogen interactions follow spatiotemporal developmental dynamics where gene expression in pathogen and host undergo crucial changes. It is of great interest to detect, quantify and localise where and when key genes are active or inactive. Here, we adapt single molecule FISH techniques to demonstrate presence and activity of mRNAs using phytomyxids in their plant and algal host from laboratory and field materials. This allowed to monitor and quantify the expression of genes from the clubroot pathogen *Plasmodiophora brassicae*, several species of its *Brassica* hosts, and of several brown algae, including the genome model *Ectocarpus siliculosus*, infected with the phytomyxid *Maullinia ectocarpii*. We show that mRNAs are localised along a spatiotemporal gradient, thus providing proof-of-concept of the usefulness of these methods. These methods are easily adaptable to any interaction between microbes and their algal or plant host, and have the potential to increase our understanding of processes underpinning complex plant-microbe interactions.

## Introduction

Thanks to a series of technological advances over the last years, it has become clear that many biological processes within and between organisms are best studied at the single cell level (e.g. Libault *et al.*, 2017; Moor & Itzkovitz, 2017). Single cell approaches provide a revolutionary toolset to study interactions, especially the complex intermingled crosstalk between biotrophic pathogens and their algal or plant hosts, when they cannot be grown outside of the host and/or are not amenable to genetic modifications. Biotrophs keep their host’s cells alive during their development and growth and therefore, have evolved a multitude of strategies to escape detection but to still obtain nutrients or water from their host (Kemen & Jones, 2012; Zeilinger *et al.*, 2016). In such interactions, timing is crucial: the pathogen first needs to escape the host defence while establishing itself, but in a subsequent step the pathogen and host communicate about nutrients to be exchanged. Hence, gene expression changes rapidly at the cellular level, and can differ between neighbouring cells (Buxbaum *et al.*, 2015). Unless they are combined with expensive or time-consuming techniques such as laser-assisted microdissection (Schuller *et al.*, 2014), widely used methods such as qPCR or RNAseq. These are not able to account for spatial and temporal heterogeneity between the cells, and therefore are impracticable to study single-cell changes at the scale of organs or macro-organisms. However, transformation or genetic manipulation remains inaccessible for a wide range of pathogens or hosts, especially those that cannot be grown in the laboratory and/or for which only very limited genetic information is available (Libault *et al.*, 2017). Furthermore, current work on plant and algal microbiomes makes increasingly clear that functional studies require to take into account complex microbial communities which include a huge diversity of "non-model” organisms, many of which are not accessible to culturing or genome amendment techniques (Egan *et al.*, 2013; Vandenkoornhuyse *et al.*, 2015; Müller *et al.*, 2016). Also, the specific developmental stages of pathogens within their hosts should be taken into account. Thus, descriptive approaches such as FISH (Fluorescence In Situ Hybridisation) have a renewed potential to start exploring these communities functionally.

Indeed, most single-cell approaches available currently lack the potential to be used routinely outside of model organisms to study the dynamic transcriptomic changes of single cells. One notable exception is the *in situ* localisation of individual mRNAs via FISH (Wang & Bodovitz, 2010; Misra *et al.*, 2014; Libault *et al.*, 2017). FISH techniques have been utilised to study gene expression patterns and the distribution of mRNA genes in human cell lines (Weibrecht *et al.*, 2013), different animal models (e.g.Trcek *et al.*, 2017), fungi and yeasts (e.g.Niessing *et al.*, 2018) and plants (Bruno *et al.*, 2011; Duncan *et al.*, 2016b; Francoz *et al.*, 2016). Recent improvements involving fluorophores, microscopic detection and resolution now allow to localise individual mRNAs of interest (Buxbaum *et al.*, 2015). Patterns of mRNA expression and the subcellular localisation of mRNAs can thus be accessed. Ultimately this leads to a better understanding of the regulatory processes behind the translation of genetic information (Chen *et al.*, 2015), without requiring genetic manipulation of the organism of interest, nor the availability of extensive genetic and molecular data. Additionally, this can be applied to field-collected samples, which opens new research lines for uncultivable organisms once an mRNA sequence of interest is available.

Phytomyxea (Rhizaria, Endomyxa) are a group of economically important plant pathogens, including e.g. *Plasmodiophora brassicae*, the clubroot pathogen, causing a loss of roughly 10 % of the world brassica crop production (Schwelm *et al.*, 2018). Additionally, ten phytomyxid species parasitize important marine primary producers, namely brown algae, seagrasses and diatoms, with essentially unknown ecological consequences (Neuhauser *et al.*, 2011; Murúa *et al.*, 2017). In brown algae, the galls formed by *Maullinia braseltonii* on the Pacific bull kelp (*Durvillea antartica*) affect the commercial value of this locally important food source. Its closely related parasite, *Maullinia ectocarpii*, infects a broad range of filamentous algae spanning at least four orders, including the genome model *Ectocarpus siliculosus* and gametophytes of the giant kelp *Macrocystis pyrifera* (Maier et al., 2000). Its availability in laboratory culture makes it a good model to start deciphering the interaction between phytomyxids and their marine hosts.

Despite their importance as plant and algal pathogens, Phytomyxea have been difficult to study, mostly because they cannot be cultured without their host and because they have a complex multi-stage life cycle (Schwelm *et al.*, 2018). This life cycle comprises two functionally different types of heterokont zoospores (primary and secondary), multinucleate plasmodia (sporangial and sporogenic), zoosporangia and resting spores. Primary zoospores (Fig. 1e) infect suitable hosts (Fig. 1f) and develop into multinucleate sporangial plasmodia (Fig. 1a) which mature (Fig. 1b, 1c) and release primary or secondary zoospores. This sporangial part of the life cycle is often restricted to few host cells. Secondary zoospores develop into multinucleate sporogenic plasmodia (Fig. 1g) which can grow to considerable size inside of the host, which in some species results in the typical hypertrophies. The sporogenic part of the life cycle ends in the formation of the resistant resting spores (Fig. 1h). Resting spores are passively released from the disintegrating host tissue and can persist for decades in the environment. Genome and transcriptome data became available only recently for *P. brassicae* (Schwelm *et al.*, 2015; Rolfe *et al.*, 2016). Yet the unavailability of genetic manipulation of the parasite forces all functional studies to be conducted on transformed plant hosts (mainly *A. thaliana*) either focussing on the host side of the response (e.g. Irani *et al.*, 2018) or by overexpressing *P. brassicae* genes in the host (Bulman *et al.*, 2019) or in other plant pathogenic fungi (Singh *et al.*, 2018). Likewise, genome resources are available for brown algae (Ye *et al.*, 2015; Cormier *et al.*, 2016), but this group yet remains inaccessible to transformation, genome editing or RNAi, to the exception of *Fucus* zygotes (Farnham *et al.*, 2013). Apart from taxonomic markers, no molecular information is currently available for *Maullinia ectocarpii* or any other marine phytomyxid nor can it be genetically modified. In this context, the prospect of monitoring gene expression of phytomyxean parasites in their host, linking the expression of selected genes of interest to specific stages of the life cycle and to specific time points in the development of the pathogen, is to get a better understanding of the interaction and the possible function of such a particular gene.

**Figure 1:**
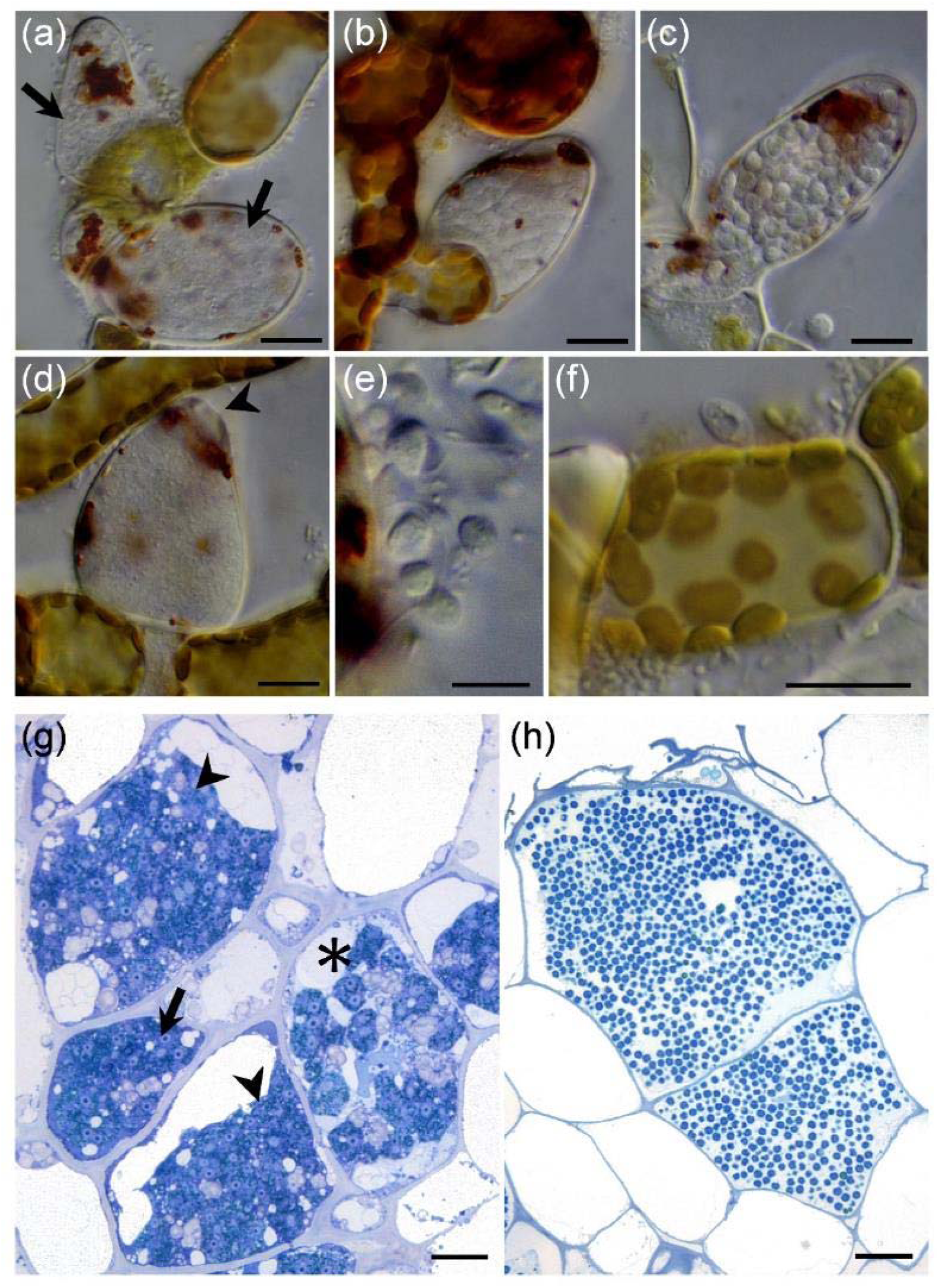
Phytomyxid Morphology. Sporangial life cycle of *Maullinia ectocarpii* (a-f) and sporogenic development of *P. brassicae* (g-h). a-f: *Maullinia ectocarpii* infecting filaments of the brown algae *Macrocystis pyrifera*. Pale cells are filled with the parasite. (a) sporangial plasmodia in enlarged algal cells (arrows). (b) sporangial plasmodium transitioning to form a zoosporangium. (c) mature zoosporangium filled with zoospores. (d) empty sporangium after the primary zoospores were released through an apical opening (arrowhead). (e) primary zoospores with two anterior flagella. (f) primary zoospore infecting the algal filament. (g, h) Chinese cabbage clubroots, cross section, methylenblue staining (g) multinucleate, sporangial plasmodia in different developmental stages. Actively growing, sporogenic plasmodia (arrow, arrowheads) and one plasmodium showing the typical lobose structure (asterisk). (h) *P. brassicae* resting spores. All resting spores inside of one host cell were formed from the same sporogenic plasmodium. Bars = 10 µm.

Here, we developed tools to monitor gene expression of intracellular pathogens and to monitor the response of their hosts (plants and brown algae) upon infection. We provide a proof-of-concept of the usefulness of single-molecule FISH to increase knowledge about the complex interactions between plants, algae and phytomyxids. For this purpose, two different approaches of mRNA localisation were evaluated: smFISH (single molecule FISH), which is based on a series of fluorescently labelled probes that tile along the mRNA of interest (Duncan *et al.*, 2016b) and RCA-FISH (Rolling Circle Amplification-FISH), which is based on *in situ* transcription of RNA followed by *in situ* -RCA signal amplification (Weibrecht *et al.*, 2013). Genes were selected on the basis of available biological background, to allow to not only test and validate FISH methods, but to also validate the feasibility and usefulness of these methods to disentangle biological information. The following *P. brassicae* genes were selected (i) the housekeeping gene *Actin1* (GenBank: AY452179.1) (Archibald & Keeling, 2004) (ii) a SABATH-type methyltransferase from *P. brassicae* (*PbBSMT*, GenBank: JN106050.1, (Ludwig-Müller *et al.*, 2015), which is able to methylate the plant defence compound salicylic acid (SA), as well as benzoic and anthranilic acids. To monitor mRNA expression and localisation in the host, the following genes were tested: (i) the *Brassica rapa* maltose excess protein 1 (*MEX1*, GenBank: XM_009109278.2) that encodes a maltose transporter, and (ii) the vanadium-dependent bromoperoxidase (*vBPO*) of *Ectocarpus siliculosus* Ec32m (Genbank: CBN73942.1), encoding a stress-inducible enzyme assumed to halogenate defensive host secondary metabolites (Leblanc *et al.*, 2015; Strittmatter *et al.*, 2016).

## MATERIAL AND METHODS

### Preparation and storage of the biological material

#### Clubroot-infected Brassica plants

##### Field sampling

Clubroot infected white cabbage (*Brassica oleracea* var. *capitata* f. *alba*), broccoli (*Brassica oleracea* var. *italica*), Chinese cabbage (*Brassica rapa ssp. pekinensis*) and kohlrabi (*Brassica olearacea* var. *gongylodes*) were collected from a commercial field (Ranggen, Austria, 47°15’24”N, 11°13’01”E). Clubroots were rinsed with tap water and treated as described below.

##### Inoculation and cultivation of plants

Chinese cabbage (*Brassica rapa* cv. *‘Granaat’*, European Clubroot Differential Set ECD-05) seeds were germinated on wet tissue paper for three days and then planted in a potting soil mixture (pH ∼ 5.7, mixing standard compost, rhododendron soil and sand 4:2:2). Plants were grown with a photoperiod of 12 h. After 12 days, plants were inoculated with 7 × 10^6^ spores of *P. brassicae*. Root galls were harvested after six to seven weeks.

##### Sample Fixation and preparation

Clubroots were cut into ca. 3 × 4 mm pieces to allow for a more homogenous fixation. Samples were transferred into Histofix 4% (phosphate-buffered formaldehyde solution, Carl Roth) where they remained for 1 - 12 h depending on sample size. Samples were used directly, or were washed in an ascending ethanol series for long-term storage (50 %, 80 %, 2x 96 %; all dilutions made with DEPC [diethyl pyrocarbonate]-treated water) and stored at −20 °C until use.

Clubroots (or clubroot pieces) were cut transversal with an RNase free razor blade by hand and washed with 1x PBS buffer (phosphate-buffered saline, 137 mM NaCl, 10 mM phosphate, 2.7 mM KCl, DEPC treated water). Making the cuts by hand posed a significantly lower risk of RNase contamination than using a cryotome (Reichert-Jung, Frigocut 2800, Suppl. Note S1).

##### Growth and maintenance of *M. ectocarpii* infected *Ectocarpus siliculosus Ec32m* and *Macrocystis pyrifera*

*Ectocarpus siliculosus* (fully sequenced genome strain Ec32m, CCAP 1310/4) was infected with *Maullinia ectocarpii* (CCAP 1538/1), using a clonal culture of *Macrocystis pyrifera* female gametophyte (CCAP 1323/1) as an intermediate host, as described by (Strittmatter *et al.*, 2016). Cultures were maintained at 15 °C with 12 h photoperiod, 20 micromol photon m^-2^ s^-1^ in artificial seawater (ASW) with half strength modified Provasoli (West & McBride, 1999). Cultures were regularly checked microscopically and samples were harvested and fixed as described above, replacing DEPC-treated water by DEPC-treated ASW. Samples were stored at −20 °C until use.

### mRNA visualisation

To prevent RNase contamination, experiments were done in RNase-free environment with RNase-free reaction mixtures. To avoid photobleaching, the samples were protected from light during and after the hybridisation of the fluorescently labelled oligonucleotides. All enzymes and reaction mixtures were kept on ice during use. Reaction tubes were incubated using a PCR cycler with heated lid to avoid evaporation or a thermal block. All incubation and amplification steps were performed in 0.2 mL PCR reaction tubes unless otherwise stated.

#### RCA-FISH (Rolling circle amplification – Fluorescence in Situ Hybridisation)

RCA-FISH is based on the *in situ* reverse-transcription of mRNA, a subsequent signal amplification using a loop-shaped DNA target probe as starting point for the RCA amplification and followed by FISH detection of the so amplified DNA (Tab. 1) (Weibrecht *et al.*, 2013). Experimentally this process can be divided into four steps: (i) Reverse transcription of target mRNAs, using a mRNA specific locked nucleic acid (LNA) primer. (ii) RNase H digestion of the RNA part of the RNA/DNA hybrid sequence, because in all subsequent steps the cDNA generated by the reverse transcription serves as template. (iii) RCA amplification: A so-called padlock probe, which forms a little loop when binding to the cDNA, serves as circular DNA template for RCA. The loop like sequence is amplified and as a part of it a sequence complementary to the detection probe. (iv) Signal detection: A standard FISH probe is used to detect the amplified signal.

**Table 1:**
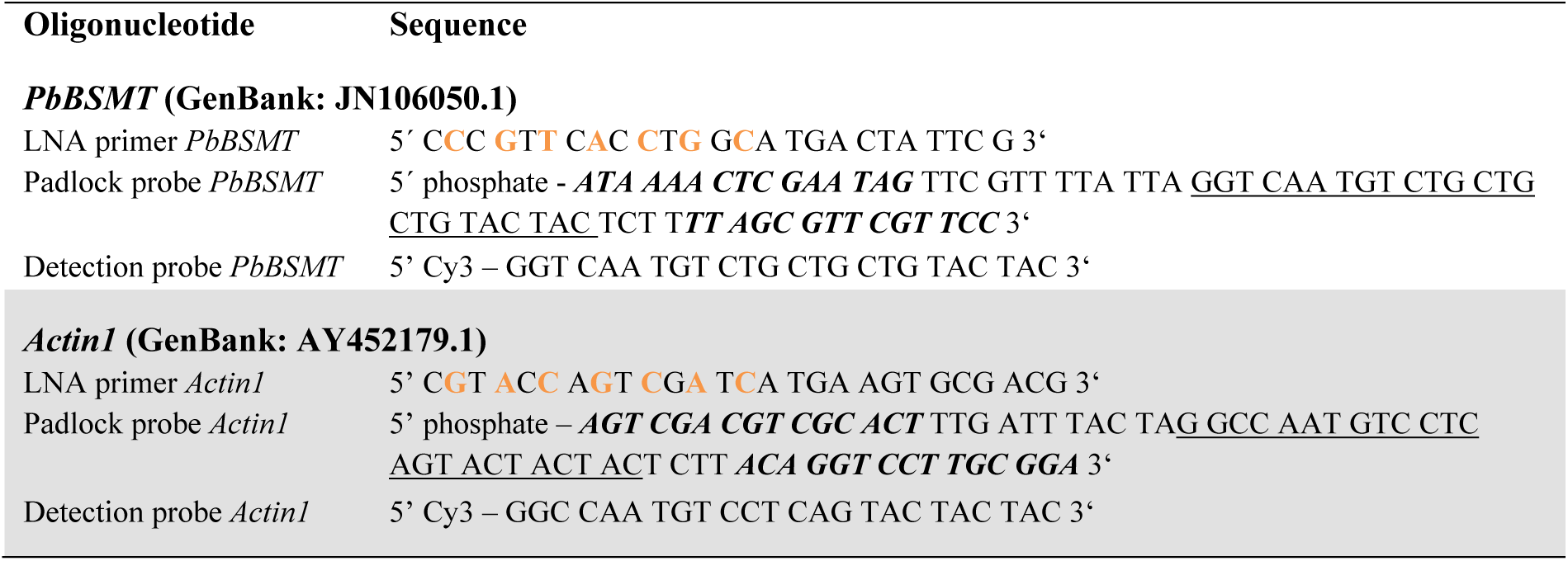
RCA-FISH probes. *PbBSMT* (*GenBank: JN106050.1*) and *Actin1 (GenBank: AY452179.1)* are genes of *P. brassicae.* LNA modified nucleotides are shown in orange, padlock probe target-specific parts are shown in italics and bold letters and the detection sequences are shown underlined.

### Primer and probe design

#### LNA-Primer

LNA primers were approximately 25 bp long and located close to the 3’ end of the mRNA. Starting from the 5’ end of the LNA primer, every second nucleotide was replaced by its LNA counterpart, in total 7 LNAs (Tab. 1).

#### Padlock Probe

About 15bp at each of the 3’ and 5’ ends of the padlock probe are complementary to the cDNA sequence. When these two regions bind to the cDNA, the central region of the probe (ca. 50 bp) forms a loop, similar to the shackle of a padlock. This central region contains a generic detection sequence (∼ 23bp) flanked by random filler sequences (Tab. 1). This loop-like structure serves as target for the Phi-Polymerase mediated RCA which is amplifying the detection sequence and consequently the signal. The padlock probe has to be phosphorylated before use. Overall the structure of the padlock probe is: 5’ phosphate – ***15bp complementary to cDNA*** – 5-15bp random filler – <u>detection sequence</u> −5-15bp random filler - ***15bp complementary to cDNA*** - 3’.

#### Detection Probe

This is a fluorescent, mono-labelled FISH probe complementary to the detection sequence included in the padlock probe.

To check the secondary structures in all three types of probes, UNAFold (https://eu.idtdna.com/UNAFold) was used. Specificity and potential off-target binding sites were checked by blasting the sequences (https://blast.ncbi.nlm.nih.gov/) against the nr database.

#### RCA-FISH experimental Method

A tick-off style, step-by-step protocol is provided in Suppl. Note S2. Modifications and notes on filamentous starting material is provided in Suppl. Note S1.

Sections from clubroot samples or algal material were transferred into the reaction tubes were they were rehydrated in PBS-T (0.05% Tween-20 in 1x PBS). Several sections may be treated in one tube, as long as the material is submerged in the reaction mix. For our samples 50 µl of reaction mix were required for consistent results. The PBS-T was removed and the reverse transcription mix (20 U µl^-1^ RevertAid H minus M-MuLV reverse transcriptase, 1 U µl^-1^ RiboLock RNase inhibitor, 1x M-MuLV RT buffer, 0.5 mM dNTP, 0.2 µg µl^-1^ BSA, 1 µM LNA primer) was added. Samples were incubated for 1 h at 37 °C in a PCR cycler. Samples were washed twice with PBS-T and were then incubated in 3.7% PFA (paraformaldehyde in 1 x PBS) for 5 min at room temperature. Samples were washed in PBS-T. This was followed by RNase H digestion, hybridisation and ligation of the padlock probe in the hybridisation mix (1 U µl^-1^ Ampligase, 0.4 U µl^-1^ RNase H, 1 U µl^-1^ RiboLock RNase inhibitor, 1x Ampligase buffer, 0.2 µg µl^-1^ BSA, 0.05 M KCl, 20% formamide, 0.1 µM padlock probe). Samples were incubated for 15 min at 45 °C and washed in PBS-T. Rolling circle amplification was done for 45 min at 37 °C (polymerase reaction mix: 1 U µl^-1^ Phi29 DNA polymerase, 1 U µl^-1^ RiboLock RNase inhibitor, 1x Phi29 buffer, 0.25 mM dNTP, 0.2 µg µl^-1^ BSA, 5% Glycerol), followed by a PBS-T washing step. The detection probe was hybridised for 10 min at 37 °C in a detection probe reaction mix (1x hybridisation mix (2x SSC saline-sodium citrate buffer (300 mM sodium chloride, 30 mM sodium citrate), 40% (vol/vol) formamide, 0.1 µM detection oligonucleotide). Samples were washed twice with PBS-T. Samples were carefully transferred and arranged on microscope glass slides, mounted with Vectashield (H-1000, Vector Laboratories) mounting medium and covered with a coverslip. The coverslips were sealed with clear nail polish and the slides were imaged using a laser scanning microscope. To exclude the possibility of false positive signals, control samples were treated without padlock probes.

#### Single molecule FISH (smFISH)

smFISH relies on a large number of mono-labelled DNA-probes complementary to the mRNA sequences of interest. The protocol described here was adapted from (Duncan *et al.*, 2016a). Probes for smFISH were designed and ordered using the Stellaris^®^ Probe Designer (http://singlemoleculefish.com, all probes labelled with Quasar570) or ordered via https://biomers.net/ (Cy3 labelled probes). A set of 48 different probes was generated, complementary to our mRNA sequences of interest (*PbBSMT* GenBank: JN106050.1, *vBPO* GenBank CBN73942.1, *MEX1* GenBank: XM_009109278.2, Suppl. Tab. S1-3). To verify the specificity of the oligonucleotides to the target sequence, the sequences were blasted against the NCBI nr database.

#### smFISH experimental method

A tick-off style step-by-step protocol is provided in Suppl. Note S3.

Samples were washed in 1x PBS buffer and were transferred to 0.2 mL PCR tubes and incubated for 1 h in 70% EtOH at room temperature. The EtOH was removed and samples were washed twice for three minutes in washing buffer (10% formamide, 2x SSC). Samples were then incubated in 50 µl of hybridisation buffer (100 mg ml ^-1^ dextran sulfate, 40% formamide in 2x SSC, 250 nM probe-mix) in a PCR cycler at 37 °C overnight with the lid heated to 100 °C to prevent evaporation. Subsequently, the hybridisation buffer was removed and the samples were rinsed twice with washing buffer, before being incubated in the washing buffer at 37 °C for 30 minutes. Nuclei were counterstained in 50 µl 4,6-diamidino-2-phenylindole (DAPI) solution (100 ng µl^-1^ in washing buffer) for 15 min at 37 °C. Samples were washed in 50 µl 2x SSC for 1 min before being equilibrated in 50 µl GLOX buffer (0.4% glucose in 10 nM Tris-HCl, 2x SSC) for 3 minutes, followed by 50 µl GLOX buffer containing enzymes (1% glucose oxidase and 1% catalase in GLOX buffer) after removal. Samples were carefully mounted using tweezers in the GLOX buffer containing enzymes. The mounted samples were sealed with nail polish and imaged as soon as possible to avoid a reduction of image quality.

To confirm RNA specificity and to evaluate autofluorescence of the samples, control samples were treated with RNase or were analysed without the addition of smFISH probes. RNase A treatment for sample sections was performed for 1 h at 37 °C (100 µg ml ^-1^) before the hybridisation step (Supplementary Note S3, between steps 4 and 5). After RNase A digestions the samples were washed twice in 10 mM HCl for 5 min and rinsed twice in 2x SSC for 5 min. These samples were then incubated with the smFISH probes as described above. Quasar570 labelled smFISH probes where used with the corresponding Stellaris buffers. Those samples were mounted in Roti-Mount FluorCare (Carl Roth).

### Image acquisition and data analyses

Images were acquired with a Leica SP5-II confocal laser scanning microscope (CLSM), using the LAS AF software (version 2.7, Leica Microsystems, Germany) and a Zeiss Cell Observer.Z1 Spinning Disk microscope using the ZEN software (Carl Zeiss Microscopy, Germany). The Leica SP5-II CLSM was equipped with a hybrid detector and a 20x (0.7 NA) or 63x (1.3 NA) objective lens and with the following lasers: 405 nm, 458 nm, 514 nm, 561 nm, 633 nm. For probes labelled with Cy3 (*Actin1, PbBSMT*, and *vBPO*) and Qu570 (*MEX1*), an excitation of 514 nm was used and the emission was detected at 550 – 585 nm. For DAPI staining, 405 nm excitation was used and emission was detected at 430 – 485 nm. The Zeiss Axio Cell Observer.Z1 was equipped with a CSU-X1 spinning disc confocal using 25x, 40x or 63x water-immersion lenses. Raw image stacks were deconvoluted with the Huygens software package (Scientific Volume Imaging, The Netherlands). Images were analysed using ImageJ (Schneider *et al.*, 2012) which was used to generate coloured overlay images of the different emissions recorded and for creating maximum projections of the z-stacks. Resizing and linear brightness and contrast adjustments were performed in GIMP 2.8.22 (www.gimp.org). The original unprocessed images will be deposited at figshare using the final numbering of the figures.

FISH signals in individual stack slices appeared dotted and therefore we used the “find maxima” process in ImageJ to segment and count these dots. With the help of the “find stack maxima” macro this process was automated for every slice in a CLSM stack (Suppl. Fig. S3, S4). CLSM settings were defined in advance to record individual signals in 2 consecutive slices to yield full signal coverage (minimum pinhole settings resulting in a slice thickness of 0.85µm; z-stepsize around 0.8 µM or below). Noise tolerance values were evaluated preliminary to best fit the fluorescence signal. Regions of interest (ROI) were marked manually (Suppl. Fig. S1, S2, S3), measured and saved. Finally, automated counts of every slice in a stack within a ROI were summarised and normalised to volume and corrected for slice number and z-stepsize. Results are presented in number of FISH signal maxima per µm^3^.

## RESULTS

Two *P. brassicae* genes were analysed using RCA-FISH: the SABATH-type methyltransferase *PbBSMT* and *Actin1*. In contrast to *Actin1*, a spatiotemporal expression pattern has already been hypothesised for *PbBSMT* since only low expression of the transcript was detected during early time points of infection and it only later increased (Ludwig-Müller *et al.*, 2015); thus we also analysed its expression using smFISH to compare the performance of both methods, qualitatively and quantitatively.

### Qualitative and quantitative comparison of smFISH and RCA-FISH

Localisation of *PbBSMT* mRNAs resulted in the same spatio-temporal pattern of dotted signals, when smFISH or RCA-FISH were used (Fig 2 a-b, see details in Suppl. Figs. S1, S2, S3, S4, S6 – S8). In z-stack maximum projections mRNA signals can appear clustered. However, single signals in individual planes appear as dots with sizes ranging from 0.2 to 0.7 µm (Suppl. Fig. S3, S4, Video S1). No signal could be detected in the controls without padlock probe (RCA FISH, Suppl. Fig S5a), after RNase treatment (smFISH, Suppl. Fig. S5b) and in uninfected plant roots (data not shown).

**Figure 2:**
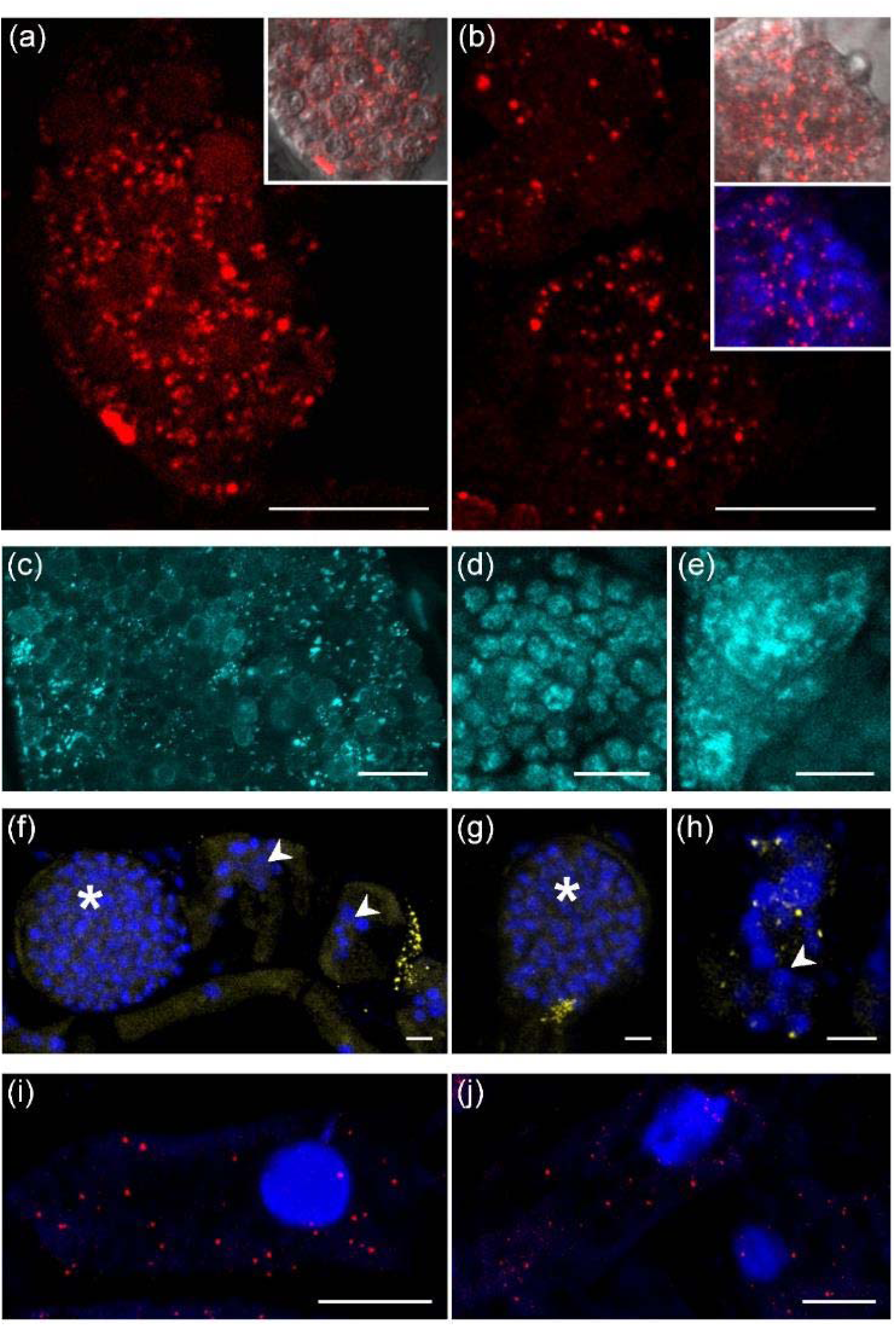
mRNA-transcript localisation in *P. brassicae* (a-e, Cy3), *Ectocarpus siliculosus* Ec32m (f-h, Cy3) and *Brassica rapa* (i-j, Qu570). (a) mRNAs of *P. brassicae PbBSMT*, RCA FISH and, (b) smFISH. The plasmodia in (a) and (b) are transitioning from the active growth phase of the plasmodium to resting spore formation, which can be recognised by the round “compartments” visible in the bright field inserts. Multiple nuclei can be seen (b, insert). (c-e): *Actin1* mRNAs of *P. brassicae* using RCA FISH (cyan). (c) during the onset of resting spore formation. (d) in the developing resting spores. (e) in actively growing sporogenic plasmodia. (f-h): Localisation of Ec32m *vBPO* mRNAs (yellow signal) using smFISH. (f) *vBPO* mRNAs close to sporangia of *M. ectocarpii*. (g, h) *vBPO* mRNAs inside of *M. ectocarpii* infected cells. arrowheads: early infection, plasmodia containing several nuclei; asterisks: later infection stage, large plasmodia with numerous parasite nuclei. (i, j): *MEX1* mRNAs in the cytosol of infected *B. rapa* root cells using smFISH (Red signals, Quasar 570). *MEX1* mRNAs are detected near the amyloplasts (black areas). Bars = 10 µm. Images of the controls of these experiments are provided in Suppl. Fig. S4.

The hypothesised spatiotemporal expression pattern of *PbBSMT* was identified with RCA-FISH (Fig. 3 g, h) and smFISH (Fig. 3i, Suppl. Fig. S8). Specifically, *PbBSMT* mRNAs started to appear in large quantities once sporogenic plasmodia started to develop into spores (Fig. 3g); the signals appeared dot-like in the individual planes, while in the maximum projection the mRNAs appeared accumulated around the developing spores. With progressing spore formation, the intensity and number of signals intensified around the outside the developing spores, both in individual layers, where most of the signals were still visible as separated dots, and in the maximum projection where mRNA signals appeared to cover the surface of the developing spores (Fig. 3 h, see 3D reconstruction in Suppl. Video S1). Subsequently, the signals became fewer and smaller until they disappeared once the spores were fully developed (Fig. 3 i). This pattern indicates that the presence of PbBSMT or its mRNA plays a role during the transition from plasmodial growth to sporogenesis in *P. brassicae*.

**Figure 3:**
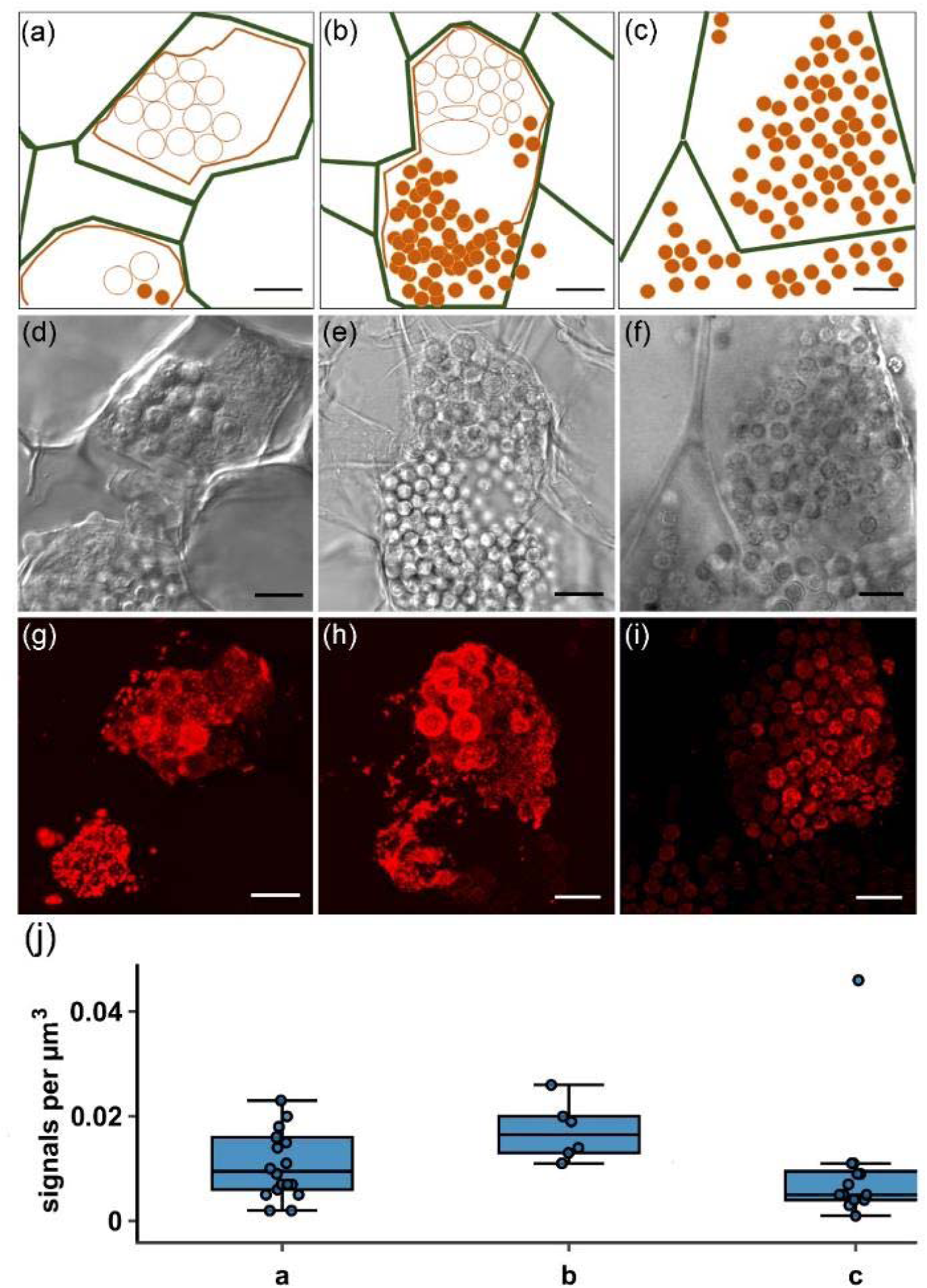
Localisation and quantification of *PbBSMT* mRNA in *P. brassicae.* a-c: diagrammatic overview of the *P. brassicae* structures seen in the brightfield images (d-e) and the FISH images (g-i). (a) The cell that is fully pictured in (a) shows a plasmodium which is at the onset of spore formation (orange polygon) inside a hypertrophied host cell (cell walls are indicated by the green lines). At the onset of spore formation round, spore like aggregations of the plasmodium become visible (orange circles). During this development stage high numbers of mRNAs can be detected (see image g) which are randomly distributed when no structures are visible in the plasmodium, but appear aggregated around the spore like structures. When the differentiation of the plasmodium progresses (b) these spore like, aggregated areas become more distinct until they have developed into individual resting spores which are not connected via the plasmodium anymore (orange dots). Signals peak around areas which already start to be recognisable as individual portions and get less in areas where resting spores are formed. Finally, the whole plasmodium has developed into resting spores (c, orange dots), that fill the entire host cell. The mRNA signals decrease further in number and brightness before disappearing completely. Bars = 10 µm. (j) Signal maxima µm-3 of smFISH *PbBSMT* signals (n=37) highlighting a peaking of *PbBSMT* mRNAs during spore formation. Labelling on the x-axes corresponds to the life cycle stages in images (a-c). No significant correlation was identified. One extreme outliner at 0.088 signals µm-3 is not displayed to improve figure layout.

In addition to the previously reported increase of *PbBSMT* expression during the later phases of the pathogen life cycle *PbBSMT* mRNAs could be detected in plasmodia that clearly were in the very early stage of secondary infection (Suppl. Fig. 5). Those plasmodia were very small in size (ca. 20 µm) and the host cells did not yet show the typical hypertrophied phenotype. This phenotype is typical for cells, which house the much larger, actively growing multinucleate plasmodia that can easily reach 100 – 150 µm in size. The young plasmodia in which the *PbBSMT* mRNAs were detected did also appear to move from one cell to the next (Suppl. Fig. S6, S7).

The number of signal maxima per µm^3^ was higher with RCA FISH (median 4,47 × 10^−02^ +/-RSD 3,01 × 10^−02^ signal maxima µm^-3^, n=9) than with smFISH (median 9,38 x× 10^−03^ +/-RSD 5,26 × 10^−02^ signal maxima µm^3^, n=37 Suppl. Tab. 4). However, when smFISH was used, the number of cells in which signals could be detected was considerably higher and more consistent across different samples. Additionally, the detection efficacy of smFISH was higher: more cells produced usable results comparing the number of positive cells per image per experiment (Suppl. Fig. S1, S2). Overall the detection efficacy of RCA FISH was lower (ca. 30% positive experiments, n = 16) compared to smFISH (ca. 45% positive experiments, n = 35). Therefore, we concluded that for detecting mRNAs in *P. brassicae* smFISH is more suitable: the reduced number of signal maxima µm^-3^ are compensated by a 5-10-fold increased number of cells with positive signals and a 15 % higher success rate. For this reason, transcripts of *Brassica oleracea* (*MEX1*) and *Ectocarpus siliculosus* (*vBPO*) were analysed with smFISH only.

### Microscopic analyses

We also compared CLSM and spinning-disk microscopy. Both methods performed very similar producing complementary mRNA detection patterns (Suppl. Fig. S7). Hand-cutting of clubroot tissue was the best and most reliable method to prepare samples for microscopy (Suppl. Note S1), which resulted in slightly uneven samples. Because of this, CLSM was the method of choice because it better allows to analyse thicker samples than wide field and spinning disk microscopy.

### Localisation of *P. brassicae Actin1* by RCA-FISH

The *Actin1* mRNA was detected in all life cycle stages of *P. brassicae* (Fig. 2c-e, Suppl. Fig S9), as bright, dot-like structures. Signals were spread evenly within cells (Fig. 2c-e) and could be visualised in individual planes (Suppl. Fig. S9 c-d), while in the maximum projection of all optical slices these dots often accumulated to larger signal hotspots (Fig 2c, Suppl. Fig. S9 e, f). In maximum projections dots were sometimes well separated (Fig 2c), while in other cells the signal was less defined appearing “blurred” and filling most of the cell without clear structural pattern (Fig 2e), indicating areas with multiple mRNA copies. The controls did not show any comparable signals, only a very weak background of autofluorescence was detected (Suppl. Fig. S5 c-e, Fig. S9 a-b).

### Localisation of host (plant and algal) transcripts during infection by phytomyxids

To evaluate whether it is possible to detect host transcripts within the same samples that host the phytomyxids, two mRNAs were chosen and analysed.

#### Brassica rapa maltose excess protein 1 (*MEX1*, smFISH)

*MEX1* encodes a transporter located in the chloroplast envelope; it is essential for the transport of maltose, the main product of starch breakdown, from the amyloplast to the cytosol (Stettler *et al.*, 2009). Starch is an important source for sugar within infected tissues to provide nutrients to *P. brassicae* as shown by the accumulation of starch grains in highly infected cells (Schuller & Ludwig-Müller, 2016). *MEX1* transcripts were detected as clear dots in the cytosol of the Brassica root cells. Dot-like signals could be observed in root cells containing amyloplasts, which were infected with *P. brassicae* (Fig. 2 i-j). The signal for *MEX1* was not present in cells filled with spores and cells that did not contain amyloplasts (Suppl. Fig. S10). Similar to the previously described genes localised detection (“dots”) could be seen in the individual optical slices and the maximum projection images indicating that mRNAs for *MEX1* are present in the whole cytosol of the plant. In both controls, the RNase treated samples and the unlabelled infected samples, no characteristic signals were detected (Suppl. Fig. S5 i-j).

#### Ectocarpus siliculosus vanadium-dependent bromoperoxidase (vBPO, *smFISH)*

mRNA detection in brown algae required adaptations of the protocol. The filamentous growth of *E. siliculosus* and of *M. pyrifera* gametophytes made it necessary to use small tufts of algal material, which are difficult to handle because of their size and shape and do not attach well to Poly L-lysine-coated slides. To reduce the risk of loosing the samples, tubes were used to incubate and wash them (see Suppl. Note S1).

*vBPO* has previously been shown to play a role in the response of *Ectocarpus siliculosus* Ec32m against pathogens (Strittmatter *et al.*, 2016) and more generally, in the responses of kelps to elicitors (Cosse *et al.*, 2009). RCA-FISH was tested without success on the algal material (data not shown), but smFISH allowed detection of *vBPO* mRNAs in Ec32m cells infected with the phytomyxid *Maullinia ectocarpii* (Fig. 2 f-h, Suppl. Fig. S11 g-i). Likewise, signals were detected in *M. ectocarpii* infected *M. pyrifera* cells (Suppl. Fig. S11 a-f). In both cases, signals were dot-like, yet much more confined - than in the plant and phytomyxids tested.

## Discussion

### mRNA localisation is a powerful tool to study pathogen – host interactions

Here, we demonstrate the feasibility of localising mRNA transcripts of phytomyxid pathogens and their plant and algal hosts, which to our knowledge is the first time this method has been used on brown algae, phytomyxids and to analyse biological interactions between two eukaryote species and in environmental samples. Localising mRNAs permits the identification and analyses of spatiotemporal pattern of mRNAs in the cytoplasm. Both methods tested (smFISH, RCA-FISH) are readily accessible once a gene of interest in biological interactions is identified. Using mRNA FISH allows to overcome important technical bottlenecks that hamper research on important non-model pathogens, plants or algae (Schwelm *et al.*, 2018). Our protocols therefore have huge potential to move our knowledge of these interactions beyond the limitations of RNAseq and organisms for which genetic amendments are possible (Buxbaum *et al.*, 2015) since the genes of interest can be studied in the environment of the direct interaction. Using clubroots from commercial fields, we directly show the applicability of the methods to investigate host-pathogen interactions in wild type populations and plants grown under natural conditions. The here presented methods will greatly increase our understanding of subcellular gene expression pattern, similar to the leap in knowledge that happened in model organisms in the coming years (Gaspar & Ephrussi, 2015; Lee *et al.*, 2016; Medioni & Besse, 2018).

FISH is – as any nucleic acid based method – very adaptable to the needs of the user. A multitude of protocols are available, many of which are variations of the two protocols tested here (reviewed by van Gijtenbeek & Kok, 2017). When selecting the most suitable method a couple of factors need to be kept in mind (Tab. 2). Generally, it can be said that the more complex a biological system, the less complex the method of choice should be: the efficacy for the detection of phytomyxid mRNAs was 15 % higher when the less complex smFISH method was used than with RCA-FISH. However, smFISH requires mRNAs which are longer than 500 bp to fit a relevant number of probes, while this size restriction does not apply to RCA-FISH. We proved here that the two contrastingly complex FISH methods can be used to generate meaningful results (Tab. 2).

**Table 2:** Advantages and disadvantages of RCA-FISH and smFISH. Prices can vary and are estimates based on Austrian prices 2019 including 20 % VAT.

|  | RCA FISH | smFISH |
| --- | --- | --- |
| **Pros** | ▪ **Signal amplification** in theory allows to monitor single copy mRNAs also in samples with a high fluorescent background.<br>▪ **Specificity:** Highly specific probes can be designed to distinguish between paralogues or SNPs .<br>▪ **No gene length limitation:** short (<500 bp) genes can be analysed.<br>▪ **Modular:** Possibility of combination with *in situ* proximity ligation assays to collect information on mRNA-protein interactions or post-translational modification (not tested in the scope of this paper). | ▪ **Signal amplification** in theory allows to monitor single copy mRNAs in samples with a high fluorescent background.<br>▪ **Complexity:** easy probe design, few steps minimising contamination risks.<br>▪ **Cell permeability:** small probes, no large functional molecules involved.<br>▪ **Cost of reagents:** most reagents needed are standard lab consumables.<br>▪ **Time:** Fast protocol, ca. 1 day from sample to image, hands on time ca 3 h. |
| **Cons** | ▪ **Complexity:** many steps increase the risk of RNase contamination or errors.<br>▪ **Cell permeability:** Involves multiple steps requiring large enzymes which need to reach the cytoplasm. This is a major drawback for use in the plant/algal system.<br>▪ **Costs of probes:** The expensive LNA- and Padlock probes need to be designed for each gene (ca. 400€ per gene for ~200 reactions).<br>▪ **Costs of reagents:** pricey reagents and enzymes are used (ca 75€ per reaction).<br>▪ **Time:** ca. 2 days from sample to image, of which there are ca 6 h hands on time. | ▪ **Specificity:** using standard FISH probes in a “string of beads” manner opens possibility of false positives for isoforms, orthologues or conserved gene families.<br>▪ **Gene length limitation:** Short mRNA sequences (< 500 bp) are not suitable for smFISH.<br>▪ **Costs of probes:** 24-48 labelled probes are needed for each gene. Some companies offer discounts on smFISH probes (ca. 500-800€ depending on the number of probes and the fluorophore, ~1000+ reactions). |

The mRNA signal pattern differed between plants, brown algae and *P. brassicae* (Fig. 2). Plant mRNAs followed the pattern that were expected based on the pioneering works on smFISH in plants (Duncan *et al.*, 2016b): mRNA signals were dot-like and distributed more or less evenly in cells and were similar in size and shape (Fig. 2i-j, Suppl. Fig. S10). The mRNA pattern in brown algae are comparable to the ones in plants, although signals were somewhat more restricted (Fig. 2f-h, Suppl. Fig. S11). The mRNA pattern observed in phytomyxids are only comparable when few mRNAs are detected in the plasmodia (e.g. Fig 2b). However, when many mRNAs are present close to each other the individual mRNAs do not resolve and are displayed as larger areas, especially in the maximum projections of the z-stacks (Fig. 2, Fig 3, Video S1). Although phytomyxid plasmodia are similar in size and shape to their host cell (Fig. 3), each plasmodium has to be interpreted as an aggregation of hundreds to thousands small cells with an individual diameter of 3-5 µm and each with its own nucleus (compare e.g. Fig 2b insert, or Fig 2 f-h). So the space which can be populated by mRNAs is much more confined than in the comparably gargantuan host cells. This is visually amplified in maximum projections of z-stacks, because one plasmodium contains more than one layer of cells (one layer per 3-5µm).. It is, however, very important to note that these technical and biological limitations are clearly counterbalanced by the amount of biological information that can be gathered.

### RNA stability and accessibility are crucial – or how do you get the protocol to work?

Here we show that both smFISH and RCA-FISH methods are suitable for *in situ* mRNA monitoring in the host and the pathogen. While analysing three totally different biological systems (plants, algae, phytomyxids) it became clear that the overall success and efficiency of the method is determined by the fixation of the mRNAs in the sample material and the permeability of the cell walls, both factors that differ across organisms. Fixation methods and the duration of storage post-fixation influenced the number of cells in which signals could be detected; however, this was different across sample types and genes so no clear maximum or minimum duration of storage could be established. Since decreasing signal intensities can be caused by RNA degradation during storage, we recommend to use the samples as soon as possible after harvesting. However, we found that samples kept in an RNase-free environment give good results after more than one year. Another reason for decreasing signals can be formaldehyde mediated covalent interlinking of mRNAs and the RNA-binding proteins which are responsible for the transport of the RNA to the site of translation (Foley *et al.*, 2017; Niessing *et al.*, 2018), or mRNA secondary structures that do not permit the binding of the probes to the sites of interest (Ding *et al.*, 2013). Because of this, we recommend not to store the samples in the formaldehyde containing fixative, but to move them to pure ethanol for long term storage. RNA-binding proteins and mRNA secondary structure can impact on the success of the detection method without the fixation bias mentioned above, because both can limit the accessibility of the target site for probes (Foley *et al.*, 2017).

Also, the permeability of cell walls and plant tissue as a whole is a constraint for FISH-based methods. In this study, RCA-FISH was not adaptable to study interactions in filamentous algae, most likely because the enzymes needed could not permeate the cell walls. Cutting of the algae without destruction of the cell arrangement is very difficult and usually results in a loss of spatial information. All tested permeabilisation efforts did not improve the signal yield. Also in plants the efficacy of the mRNA detection was lower in RCA-FISH, as signals were only visible in cells which were cut open, but never in cells with an intact cell wall. The cooperatively small probes used for smFISH could be used on all samples without additional permeabilisation steps.

### mRNA localisation sheds light on phytomyxid biology

In this study, we determined the expression and localisation pattern of three pathogen genes, one plant and one brown algal gene. All mRNAs chosen had a putative function assigned to pre-existing information on the expected expression pattern. However, the single cell resolution of our experiments resulted in information, which is already improving our understanding of the biological interaction. These findings also showcase the potential gain of using mRNA FISH to advance our knowledge on plant pathogen interactions beyond the state of the art.

Biologically most interesting was the expression pattern of *PbBSMT*, a SABATH-type methyltransferase produced by *P. brassicae*. This methyltransferase has structural similarities to plant methyltransferases and is able to methylate SA. Previous qPCR analysis of its expression pattern showed, that its expression during clubroot development is highest when the concentration of SA in the roots peaks, which is the reason why a role in disease development of this gene has been discussed (Ludwig-Müller *et al.*, 2015; Bulman *et al.*, 2019). In our *in situ* experiments, *PbBSMT* mRNAs started to appear in small developing plasmodia (Suppl. Fig S6, S7). Small amounts of *PbBSMT* have been detected previously during early infection in EST (expressed sequence tag) libraries (Bulman *et al.*, 2006) and callus culture transcriptome analyses (Bulman *et al.*, 2011). We observed mRNAs of *PbBSMT* in small plasmodia, which appear to move from cell to cell (Suppl. Fig. S7). There is anecdotal evidence that plasmodia can move from cell to cell using cell wall breaks or plasmodesmata (Mühlenberg *et al.*, 2003; Donald *et al.*, 2008; Riascos *et al.*, 2011), but our results are the first to show that this movement is linked to the expression of a putative effector that alters the host defence response.

Using FISH we clearly demonstrate, that *PbBSMT* mRNAs start to increase when *P. brassicae* transitions from plasmodial growth to resting spore formation. The detected mRNAs peaked when young, immature resting spores became recognisable (Fig. 3). These results again confirm findings from previous studies (Ludwig-Müller *et al.*, 2015; Bulman *et al.*, 2019), but our results are the first to pinpoint these changes to specific life cycle stages. Notably *PbBSMT* mRNAs accumulate around the developing resting spores in the sporogenic plasmodia (Fig. 3, Suppl. Video 1). This accumulation of *PbBSMT* mRNAs around the developing resting spores is striking, because during spore formation chitin is produced (Cavalier-Smith & Chao, 2003; Schwelm *et al.*, 2015), which is one of the best studied elicitors of plant defence and induces (amongst others) SA production (Fesel & Zuccaro, 2016). The results presented here therefore reinforce the previously established hypothesis that *PbBSMT* is produced to inactivate SA produced by the host in response to chitin (Ludwig-Müller *et al.*, 2015; Schwelm *et al.*, 2015). Howerver, methylsalicylate (MeSA) is a more volatile and membrane-permeable form of SA and is either excreted via the leaves or an inducer of systemic responses in plants (Vlot *et al.*, 2017). MeSA was specifically emitted from Arabidopsis leaves infected with *P. brassicae* (Ludwig-Müller *et al.*, 2015). Combined with findings from previous studies (Ludwig-Müller *et al.*, 2015; Bulman *et al.*, 2019), our observations strongly support the role in host defence suppression of *PbBSMT*, making it a very interesting target for future studies.

Therefore, we could confirm a direct interaction and interference of phytomyxids on the transcriptional status of infected cells using smFISH. Roots of brassicas infected with *P. brassicae* show a marked accumulation of starch (Ludwig-Müller *et al.*, 2009; Schuller & Ludwig-Müller, 2016). *MEX1*, a gene coding for a maltose transporter essential for transporting maltose from the amyloplast to the cytosol (Stettler *et al.*, 2009). With smFISH we could localise *MEX1* mRNAs in plant cells containing amyloplasts and plasmodia of *P. brassicae* (Suppl. Fig. S10). The presence of mRNAs is an indicator of ongoing synthesis of a host protein, because of the short half live of mRNAs in vivo (Merchante *et al.*, 2017). Therefore, it is likely that the MEX1 protein is activated by growing *P. brassicae* plasmodia to mediate energy supply from the host to the pathogen.

The brown algal *vBPO* has previously been linked to stress response (Leblanc *et al.*, 2015; Strittmatter *et al.*, 2016). Here we could confirm that *vBPO* mRNAs are located in *E. siliculosus* and *M. pyrifera* cells, which show early infections with *Maullinia ectocarpii*, while in cells where the infection has advanced to sporangial development no signals were identified. This confirms that brown algae, like plants, show a localised stress response to plasmodiophorid pathogens. Whether or not this type of stress response is a general pattern in brown algae or if this is specific for infections with phytomyxids was beyond the scope of this study.

## Supporting information

Supplemtn image analysis

## Acknowledgements

JB and SN were funded by the Austrian Science Fund (FWF): grant Y801-B16 (START-grant). CG has received funding from the European Union’s Horizon 2020 research and innovation programme under the Marie Sklodowska-Curie grant agreement No 642575, and from the UK NERC under the grant agreement GlobalSeaweed (NE/L013223/1). We thank Stefan Ciaghi for providing the transcriptomic data for *MEX1*. The authors want to thank Martin Kirchmair, Arne Schwelm, Mohammad Etemadi and Stefan Ciaghi for useful discussions.

## Author Contribution and References

JB and SN designed the research, collected, analysed and interpreted data and wrote the manuscript with feedback from all co-authors; JB performed the research; CG provided algal material and algal data; AMS analysed the images. All authors read and contributed to the final MS.

