## Supplementary Figures for "Biotrophic interactions disentangled: *In situ* localisation of mRNAs to decipher plant and algal pathogen – host interactions at the single cell level"

### ELECTRONIC SUPPLEMENTARY MATERIAL

Julia Badstöber[1], Claire M. M. Gachon[2], Jutta Ludwig-Müller[3], Adolf M. Sandbichler[4], Sigrid Neuhauser[1]
[1]Institute of Microbiology, University of Innsbruck, A-6020 Innsbruck, Austria
[2]The Scottish Association for Marine Science, Scottish Marine Institute, Oban PA37 1QA, UK
3Institute of Botany, Technische Universität Dresden, D-01217 Dresden, Germany
[4]Institute of Zoology, University of Innsbruck, A-6020 Innsbruck, Austria

Author for correspondence:
*Sigrid Neuhauser*
**

**Supplementary Video**

Video S1: 3D reconstruction of *PbBSMT* mRNA signals – see file Video S1 Additional file

**Supplementary Notes**

*Note S1:* RCA-FISH: modifications for filamentous starting material p. 2
*Notes S2*: Step-by-step Protocol for RCA-FISH p. 3
*Notes S3:* Step-by-step protocol for smFISH p. 6

**Supplementary Figures**

Fig. S1: RCA-FISH images used for signal quantification (*PbBSMT*, *P. brassicae*) p. 8
Fig. S2: smFISH images used for signal quantification (*PbBSMT*, *P. brassicae*) p. 9
Fig. S3: Comparison of signals in maximum projection and individual z-stacks. p. 10
Fig. S4: Comparison of signals in maximum projection and individual z-stacks. p. 11
Fig. S5: Control experiments accompanying Fig. 1 main-text. p. 12
Fig. S6: *P. brassicae, PbBSMT* in young secondary plasmodia, smFISH p. 13
Fig. S7: comparison of spinning disk and confocal laser scanning microscopy p. 14
Fig. S8: *Plasmodiophora brassicae, PbBSMT*, smFISH p. 15
Fig. S9: *Plasmodiophora brassicae, Actin1*, RCA-FISH p. 16
Fig S10: *Brassica napa*, *MEX1*, smFISH p. 17
Fig. S11: *Ectocarpus siliculosus* Ec32m, *vBPO*, smFISH p. 18
Fig. S12: Effect of formamide concentrations on *PbBSMT* detection, smFISH p. 19

**Supplementary Tables**

Tab. S1: smFISH probes Ec32m *vBPO* p. 20
Tab. S2: smFISH probes *P. brassicae PbBSMT* p. 21
Tab. S3: smFISH probes *B. rapa MEX1* p. 22
Tab. S4: z-stack thickness and number of planes p. 23
Tab. S5: Signal counts – see excel file Suppl. Tab S5 Additional file

#### Note S1: RCA-FISH: modifications for filamentous starting material algal cultures

For the *Maullinia-Ecotacarpus* pathosystem different approaches were tested, but smFISH was superior. This is the reason why we did not optimize the RCA protocol to an “application ready” stage. However, some of the approaches tested might be useful for other biological systems and are outlined below.

##### Modification 1: Cutting procedure

Cutting with RNase free razor blades and a cryotome (Reichert-Jung, Frigocut 2800) were tested. For clubroot samples using razorblades gave clearly better results than cuts produced by a cryotome. We hypothesize this is due to the reduced risk of RNase contamination as less equipment needs to be cleared from RNase. If thinner, serial or very even sections are needed for an experiment, the setup needs to be optimized, and other cutting devices might be more suitable than the one tested.

Filamentous starting material such as algae, fungi or oomycetes cannot easily be cut without losing spatial information with both methods. If filamentous material needs to be cut, caution should be taken to have the cell layers as aligned as possible to maximize the amount of neighboring cells that are cut at the same level.

##### Modification 2: Chemical and enzymatic permeabilisation

To allow the enzymes used for RCA-FISH to enter the cells of interest chemical and/or enzymatic permeabilisation is an alternative. A number of classical chemical approaches (Triton X, cellulase/pectinase and proteinase K) were tested without improving yield for our samples. We also tested Glucanex (Sigma-Aldrich, L1412) a mixture of β-glucanase, cellulose, protease and chitinase activities) again without improved yields. All the treatments described above did not have negative impact on the results.

##### Modification 3: Hybridization chambers

Use Poly L-lysine slides combined with self-adhesive hybridization chambers (e.g. Secure Seal chambers, Grace Bio-Labs) for all hybridization and washing steps instead of 1.5 mL tubes. The reaction volume should be adjusted to fill the chamber completely. Seal the chambers during each incubation step with self-adhesive PCR film to avoid evaporation.

The hybridization chambers need to carefully removed with tweezers after the last washing step (step 19 in the protocol given in Note S1). Before mounting the samples should be allowed to air dry for a few seconds before mounting and sealing them.

**Notes S2: Step-by-step protocol for RCA FISH on clubroot samples**

This protocol is based on the method described by Weibrecht et. al. 2013. "*In situ detection of individual mRNA molecules and protein complexes or post-translational modifications using padlock probes combined with the in situ proximity ligation assay.*" Nat Protoc 8(2): 355-372.

---

NB: This protocol has been optimized for use with clubroot tissue. For modifications suggested for filamentous starting material such as algae of fungi see Notes S2.

---

☐ **Sample Fixation (1 h – 2 days)**

Depending on the starting material and the time the material should be stored two approaches were used:

A) Material for immediate use:
   - Incubate clubroots in Histofix 4% (phosphate-buffered formaldehyde solution (PFA), Carl Roth, P087.6) for 1-12 h depending on the size of the clubs and store them at 4 °C for up to 7 days.

B) Material for long term storage:
   - Cut clubroots into smaller pieces and incubate in Histofix 4% (phosphate-buffered formaldehyde solution (PFA), Carl Roth, P087.6) for 1 h.
   - Incubate the samples in 50% EtOH for at least 1 h, preferably overnight.
   - Incubate samples in 80% EtOH for 1 h. Repeat this step.
   - Incubate samples in 96% EtOH preferably overnight. Repeat this step.
   - The samples can be stored at – 80 °C or - 20 °C in 96% EtOH.

---

CAUTION: PFA is harmful. All contact with skin, eyes and clothes should be avoided.

---

☐ **Sample preparation (cutting, 10 min – 1h)**

1. Cut the biological material with RNase-Away (Carl Roth, A998.1) treated razor blades into suitable sections, and wash it with 1x DEPC treated PBS buffer (Thermo Scientific, AM9624). For other cutting techniques or chemical and enzymatic permeabilisation steps see Note S2.

☐ **Reverse transcription (1 h 30 min)**

2. Rehydrate the sample sections in PBS-T (0.05% Tween-20 in 1x PBS) for a few minutes (e.g. while preparing the reaction mix described in step 3) in a 0.5 mL PCR tube.
3. Mix the "Reverse-transcription mix" in a 1.5 mL tube according to the list below.

| **Reverse transcription mix** | **Per reaction** | |
|---|---|---|
| RevertAid H minus M-MuLV reverse transcriptase (200 U $\mu l^{-1}$, Thermo Scientific, EP0451) | 5 | µl |
| RiboLock RNase inhibitor (40 U $\mu l^{-1}$, Thermo Scientific, EO0381) | 1.25 | µl |
| M-MuLV RT buffer, 5× (included in Thermo Scientific, EP0451) | 20 | µl |
| dNTP (10 mM, Thermo Scientific, R0191) | 2.5 | µl |
| BSA (10 µg $\mu l^{-1}$, Carl Roth, 8895.1) | 1 | µl |
| LNA primer (10 µM, Eurogentec) | 5 | µl |
| DEPC-$H_2O$ | 20.25 | µl |
| Total | 55 | µl |

4. Remove the PBS-T from the sample and add the reverse transcription mix.
5. Incubate the samples at 37 °C for 60 min.

6. Remove the reverse transcription mixture and wash twice with PBS-T.

PAUSEPOINT: The samples can be stored in PBS-T overnight at 4 °C

7. Fix cells in freshly prepared 3.7% PFA in DEPC-PBS for 5 min at RT.
8. Wash the cells and tissue samples in PBS-T.

PAUSEPOINT: The samples can be stored in PBS-T overnight at 4 °C

##### ☐ RNase H digestion, hybridization and ligation with ampligase (30 min)

9. Mix the "Ligation Mix" reagents according to the list below in a 1.5 mL reaction tube.

| **Ligation Mix** | **Per reaction** |
|---|---|
| Ampligase (5 U µl$^{-1}$, Lucigen, A0102K) | 10 µl |
| RNase H (5 U µl$^{-1}$, Thermo Scientific, EN0201) | 4 µl |
| RiboLock RNase inhibitor (40 U µl$^{-1}$ Thermo Scientific, EO0381) | 1.25 µl |
| Ampligase buffer, 10× (included in Lucigen, A0102K) | 5 µl |
| BSA (10 µg µl$^{-1}$, Carl Roth, 8895.1) | 1 µl |
| KCl, 1 M | 2.5 µl |
| Formamide (≥99,5%, Sigma-Aldrich F9037) | 10 µl |
| Padlock probe, (2 µM, Eurogentec) | 2.5 µl |
| DEPC-H2O | 13.75 µl |
| Total | 50 µl |

10. Remove the PBS-T from the tube containing the samples and add the ligation mix.
11. Incubate the samples for 15 min at 45 °C.
12. Wash the samples in PBS-T.

PAUSEPOINT The samples can be stored in PBS-T overnight at 4 °C

##### ☐ Rolling circle amplification (1 h)

13. Mix the "RCA-Mix" reagents in a 1.5 mL tube following the list below.

| **RCA Mix** | **Per reaction** |
|---|---|
| Phi29 DNA polymerase (10 U µl$^{-1}$, Thermo Scientific, EP0091) | 5 µl |
| RiboLock RNase inhibitor (40 U µl$^{-1}$ Thermo Scientific, EO0381) | 1.25 µl |
| Phi29 buffer 10× (included in Thermo Scientific, EP0091) | 5 µl |
| dNTP (10 mM, Thermo Scientific, R0191) | 1.25 µl |
| BSA (10 µg µl$^{-1}$ Carl Roth, 8895.1) | 1 µl |
| Glycerol 50% (vol/vol) (Sigma-Aldrich, G5516) | 5 µl |
| DEPC-H2O | 31.5 µl |
| Total | 50 µl |

14. Remove the PBS-T from the tube containing the sample and add the RCA mix.
15. Incubate the samples for 45 min at 37 °C.
16. Wash the samples in PBS-T.

PAUSEPOINT The samples can be stored in PBS-T overnight at 4 °C.

☐ **Detection probe hybridization (30 min)**

---

NB: From this step onward, the samples should be protected from light in order to avoid weakening of the fluorophores!

---

17. Prepare the "detection mix" reagents according to the list below.

| **Detection Mix** | **Per reaction** |
|---|---|
| Hybridization mix, 2× (contains: 4x SSC (Saline-sodium citrate buffer) Thermo Scientific, AM9763 and 40% (vol/vol) formamide (≥99,5%, Sigma-Aldrich F9037)) | 25 µl |
| Detection oligonucleotide, 10 µM (Biomers) | 0.5 µl |
| DEPC-H2O | 24.5 µl |
| Total | 50 µl |

18. Remove the PBS-T from the tube containing the sample and add the detection mix.
19. Incubate the samples at 37 °C for 10 min.
20. Wash the samples in PBS-T.

---

NB: When additional structures should be stained (e.g. calcofluorwhite) this can be performed at this stage.

---

21. Carefully transfer the samples from the 1.5 mL tube onto microscope slides. Take care to evenly spread the samples.
22. Mount the samples in a slow fade medium (e.g. Vecatshield), add a coverslip and seal.
23. Image slides with a suitable high resolution microscope.

**Notes S3: "Step-by-step" protocol for smFISH on clubroot samples**

This protocol is based on the method described by Duncan et. Al. 2016. "*A method for detecting single mRNA molecules in Arabidopsis thaliana.*" Plant Methods 12: 13.

☐ **Sample preparation (cutting, 10 min – 1h)**

Cut the fresh root galls with RNase-Away (Carl Roth, A998.1) treated razor blades into suitable sections. For modifications suggested for filamentous starting material such as algae of fungi see Notes S2.

☐ **Sample Fixation (1 h – 3 h)**

Depending on the size and thickness of the starting material, incubation time has to be adjusted.

1. Incubate clubroot sample sections in Histofix 4% (phosphate-buffered formaldehyde solution (PFA), Carl Roth, P087.6) for 1 - 3 h depending on the size of the sample sections at room temperature (RT).

---

CAUTION: PFA is harmful. All contact with skin, eyes and clothes should be avoided.

---

☐ **Sample permeabilisation (1.5 h)**

2. Wash the samples in 1x PBS buffer (phosphate-buffered saline, 137 mM NaCl, 10 mM phosphate, 2.7 mM KCl, DEPC treated water) and transfer them to 0.2 mL PCR-reaction tubes.
3. Incubate the sections for 1 h in 70% EtOH at RT.
4. Remove the EtOH and wash them twice for 3 min in washing buffer (10% formamide, 2x SSC) (20x saline-sodium citrate buffer (SSC), Thermo Scientific, AM9763). Remove the washing buffer.

☐ **Hybridization (overnight)**

The formamide concentration must be adjusted, as well it is dependent from probe composition. Experiments were performed, using different formamide concentrations (Suppl. Fig. S11). For these experiments, 40% formamide delivered the best results, therefor we decided to use this formamide concentration.

5. Incubate the samples in 50 µl of hybridization buffer containing probes (250 nM) overnight at 37 °C in a PCR cycler (lid heat 100 °C to prevent evaporation).

| **Hybridization buffer (10 mL stock)** | |
|---|---|
| Dextran sulfate (Sigma, Res2029D-A7) | 1 g |
| Nuclease free 20x SCC (Thermo Scientific, AM9763) | 1 mL |
| Formamide | 4 mL |
| Nuclease-free water | Fill up to 10 ml |

6. Remove the hybridization buffer and wash the samples twice with washing buffer.
7. Incubate the samples for 30 min at 37 °C in washing buffer.

☐ **DAPI staining and sample mounting**

8. Stain in 50 µl DAPI solution (100 ng $\mu l^{-1}$ washing buffer) for 15 min at 37 °C.
9. Wash the samples in 50 µl 2x SCC for 1 min.
10. Equilibrate the sections in 50 µl GLOX buffer for 3 min. (GLOX buffer is not compatible with FAM labelling. Alternatively, vectashield mounting medium (H-1000, Vector Laboratories) or Roti-Mount FluorCare (Carl Roth) could be used.)

**GLOX buffer (1 mL stock)**

| | |
|---|---|
| 10% glucose in nuclease free water | 40 µl |
| 1 M Tris-HCl (Thermo Scientific, AM9855G) | 10 µl |
| 20x SCC (Thermo Scientific, AM9763) | 100 µl |
| Nuclease-free water | 850 µl |

11. Remove the GLOX buffer and add GLOX buffer containing enzymes.

**GLOX buffer containing enzymes (100 µl)**

| | |
|---|---|
| Glucose oxidase (Sigma, G0543) | 1 µl |
| Catalase suspension (Sigma, C3155) | 1 µl |
| GLOX buffer (without enzymes) | 100 µl |

12. Mount sample sections carefully with tweezers on glass slides (in GLOX buffer containing enzymes) and add a coverslip.
    (For samples labelled with FAM, Roti-Mount Fluor Care mounting medium could be used)
13. Seal the coverslips with clear nail polish.
14. Image the slides with a high resolution microscope.

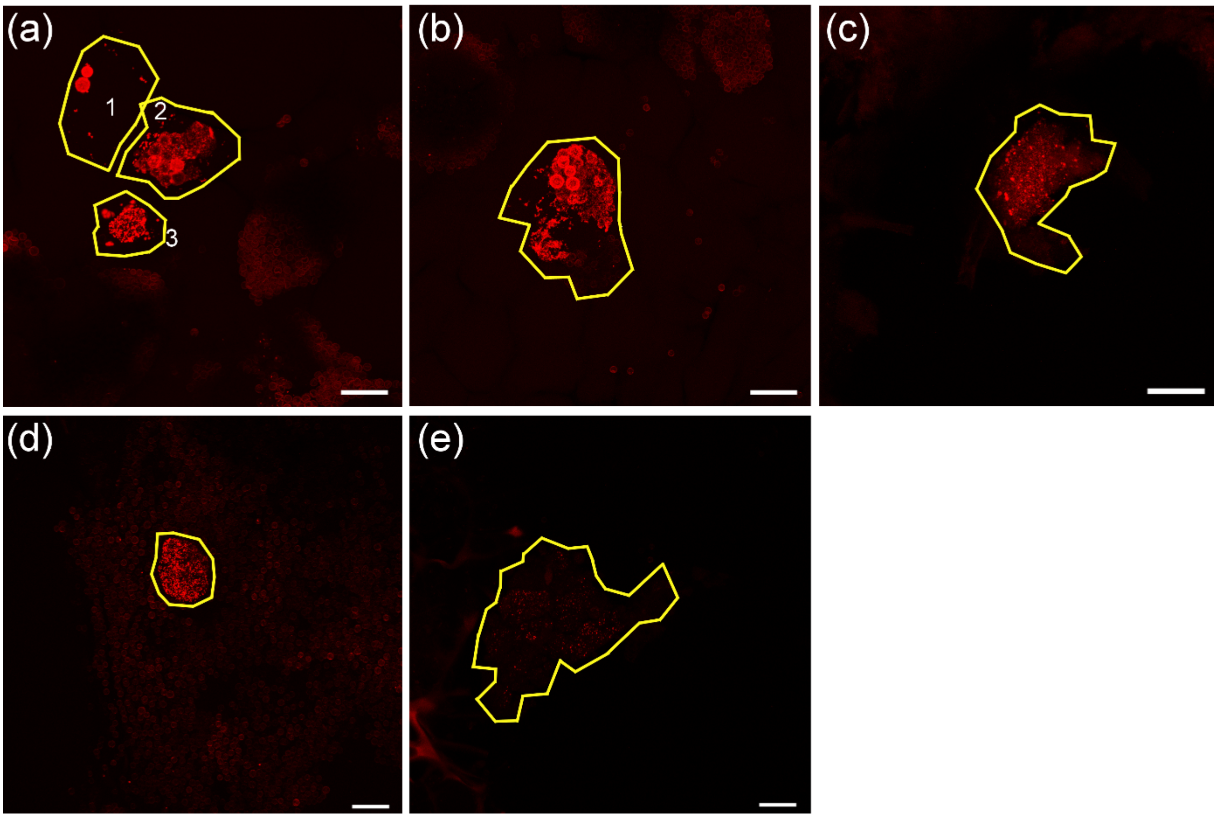


**Figure S1: Selection and localisation of cells for *PbBSMT* mRNA quantification (displayed in red pseudocolour), RCA FISH, *P. brassicae*.** Regions of interest (ROIs) were defined based on the host cell walls and the margins of selected ROIs is highlighted by yellow lines. Each ROI was analysed individually (see Supplementary Tab. S5). Each of the ROIs was assigned as developmental stage as described in Fig. 2a-2c.

Plasmodia at the onset of resting spore formation, the plasmodium structure is still visible but starts to partition into spores, corresponding to Fig. 3a: cells a2, c.

Plasmodia which have nearly fully undergone transformation into resting spores. Most of the multinucleate plasmodium has partitioned into single cell portions, some fully developed resting spores are already visible, corresponding to Fig. 3b: cells a3, b, d.

More or less mature resting spores. The plasmodium has fully undergone transition into resting spores which already start to show the characteristic thick cell wall, corresponding to Fig. 3c: cells a1, e.

All images are presented as maximum projection. Cross sections of Chinese cabbage roots (galls), infected with *P. brassicae*. Bars = 20 µm

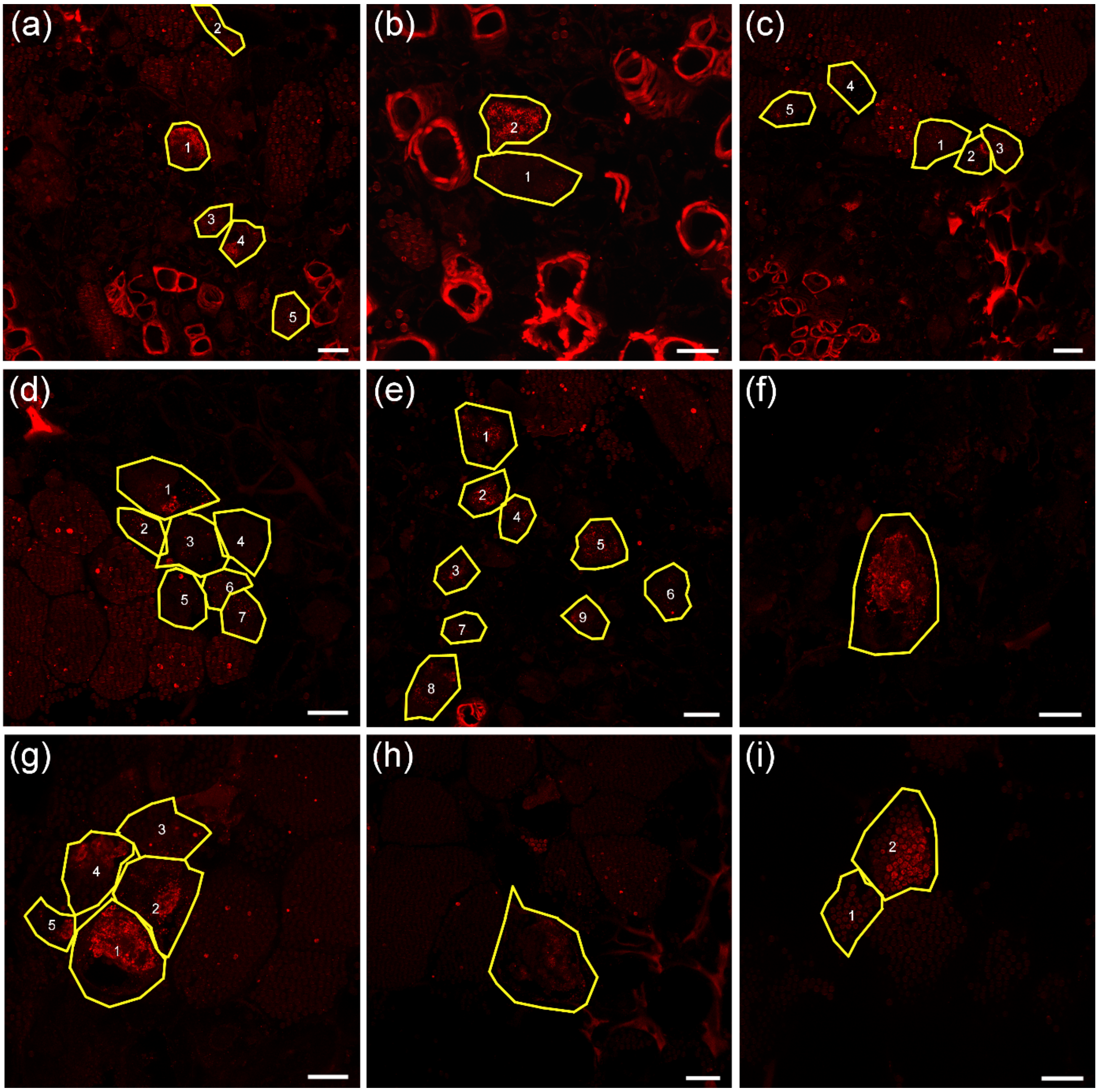


**Figure S2: Selection and localisation of cells for *PbBSMT* mRNA quantification (displayed in red pseudocolour), smFISH, *P. brassicae*.** Regions of interest (ROIs) were defined based on the host cell walls encircled by yellow lines. Each ROI was analysed individually. Each of the ROIs was assigned a developmental stage as described in Fig. 2a-2c.

Plasmodia at the onset of resting spore formation. Plasmodium is still visible but starts to partition into spores, corresponding to Fig. 3a: a1-4, b1-2, c2-3, d1-3, d5-7, e5, f, g2, h.

Plasmodia which have nearly fully undergone transformation into resting spores. Most of the multinucleate plasmodium has partitioned into single cell portions, some fully developed resting spores are already visible, corresponding to Fig. 3b: e1-2, g1, g4-5, i2.

More or less mature resting spores. Plasmodium has fully undergone transition into resting spores, which already start to show the characteristic thick cell wall, corresponding to Fig. 3c: a5, c1, c4-5, d4, e3-4, e6-9, g3, i1.

Pictures are presented as maximum projection. Cross sections of Chinese cabbage roots (galls), infected with *P. brassicae.* Bars = 20 µm.

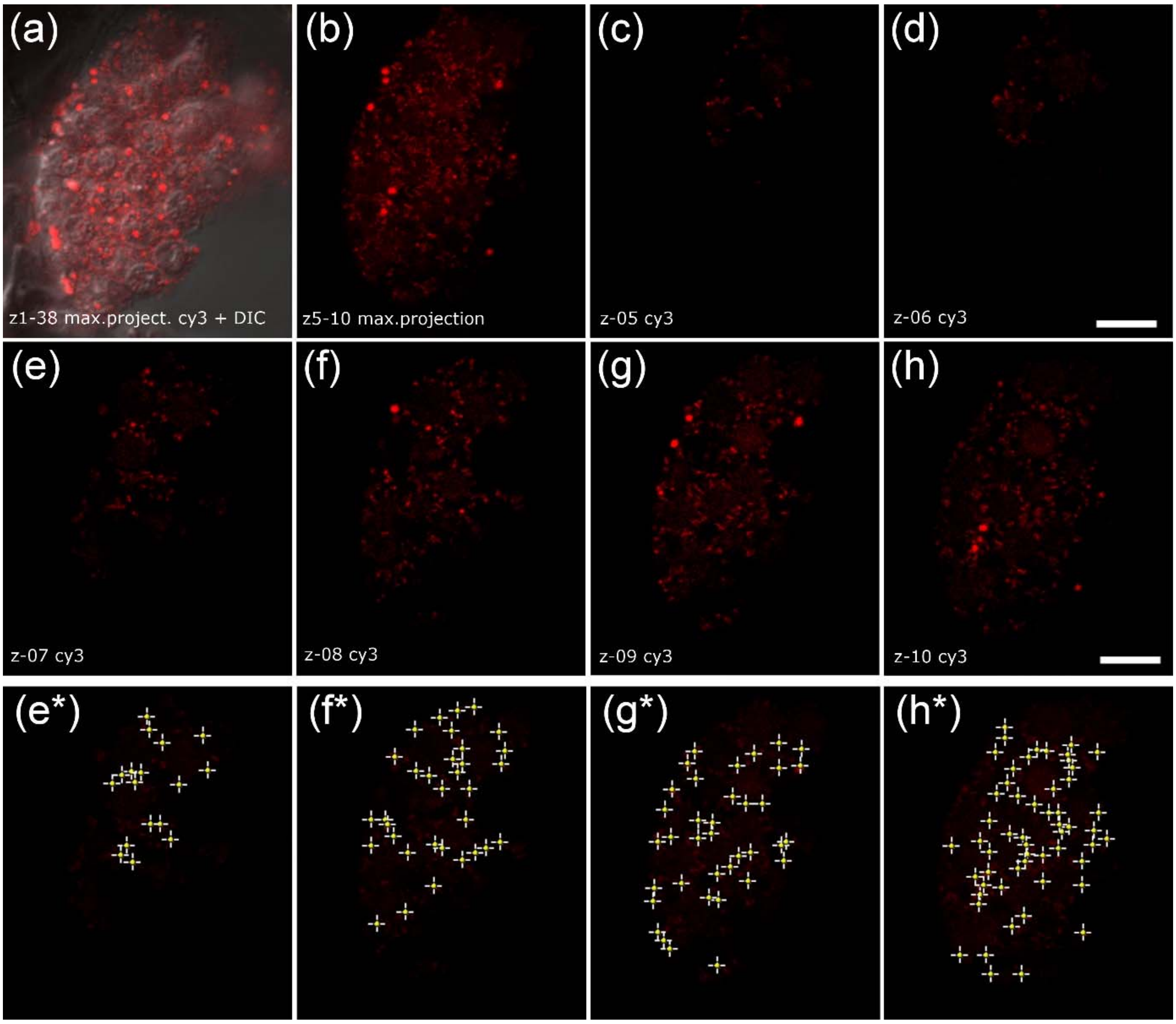


**Figure S3: Demonstration of the formation of the apparent signal clustering in z-stack maximum projections and comparison to single z-plane images, and visualized signal counting.** (a) – (h) RCA-FISH of *PbBSMT* (Cy3, red) in Chinese cabbage clubroot cross section, CLSM. (a) z-stack maximum projection of 38 individual planes with a slice thickness of 0,8 µm, distance between top and bottom z- stack 30,4 µm. Overlay of cy3 channel with brightfield image. (b) maximum projection of images z05-10; c-h: examples of individual z-slices. (c) z-05, margin of plasmodium, only very few, small signals are visible. Individual dot-like red signals are close to each other. (d) – (h) subsequent z-planes (z 07-09) (h) z-10, centre of the plasmodium. (e*) – (h*): visualized signal counting with ImageJ, of images (e) – (h). Images a-h show a plasmodium at the onset of resting spore formation when the multinucleate plasmodium is still visible, but starts to partition into spores (images (a) – (h) correspond to Fig. 2a in the maintext). Bars = 5 µm

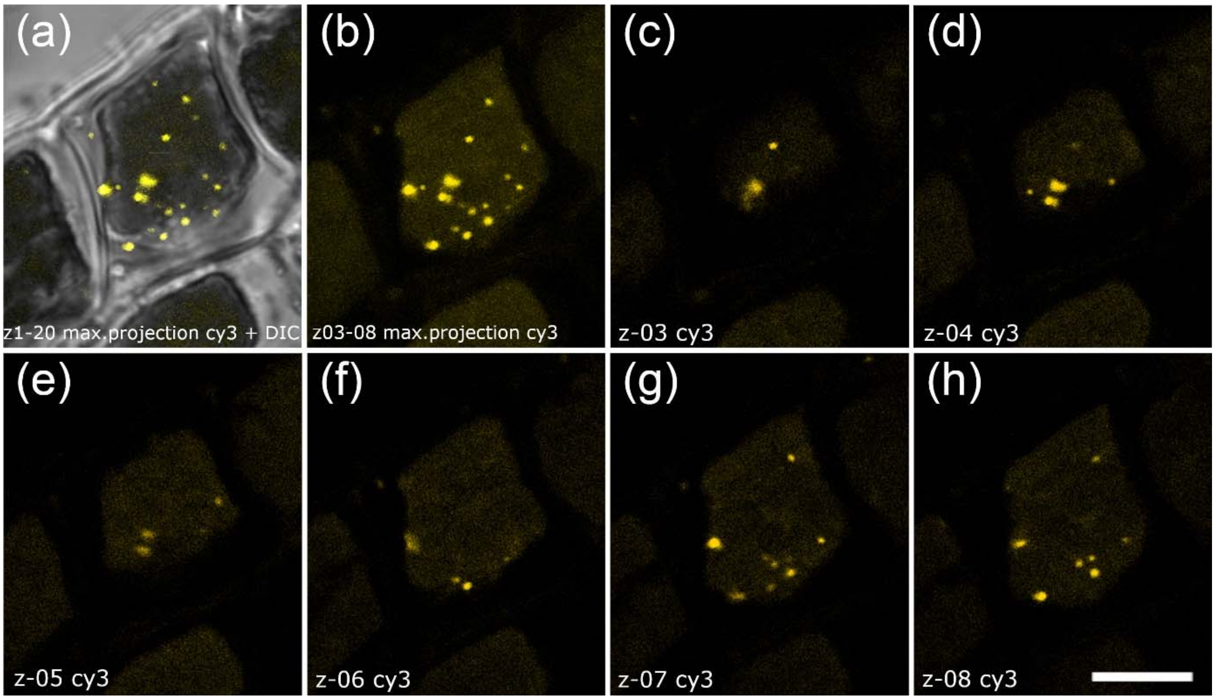


**Figure S4: Demonstration of the formation of the apparent signal clustering in z-stack maximum projections and comparison to single z-plane images, and visualized signal counting.** (a) – (h): smFISH of *vBPO* (Cy3, yellow) in *Maullinia ectocarpii* infected *Macrocystis pyrifera,* CLSM. (a) z-stack maximum projection of 20 individual planes with a slice thickness of 0,8 µm, distance between top and bottom z- stack 15,15 µm. Overlay of cy3 channel with brightfield image. (b) maximum projection of images z03-08. (c) – (h) subsequent z-planes (z03-Z08). Individual dot-like yellow signals are close to each other. (a) – (h) correspond to Fig. S11 in the supplementary information Images (a) – (h) show an infected brown algal cell. Bars = 5 µm

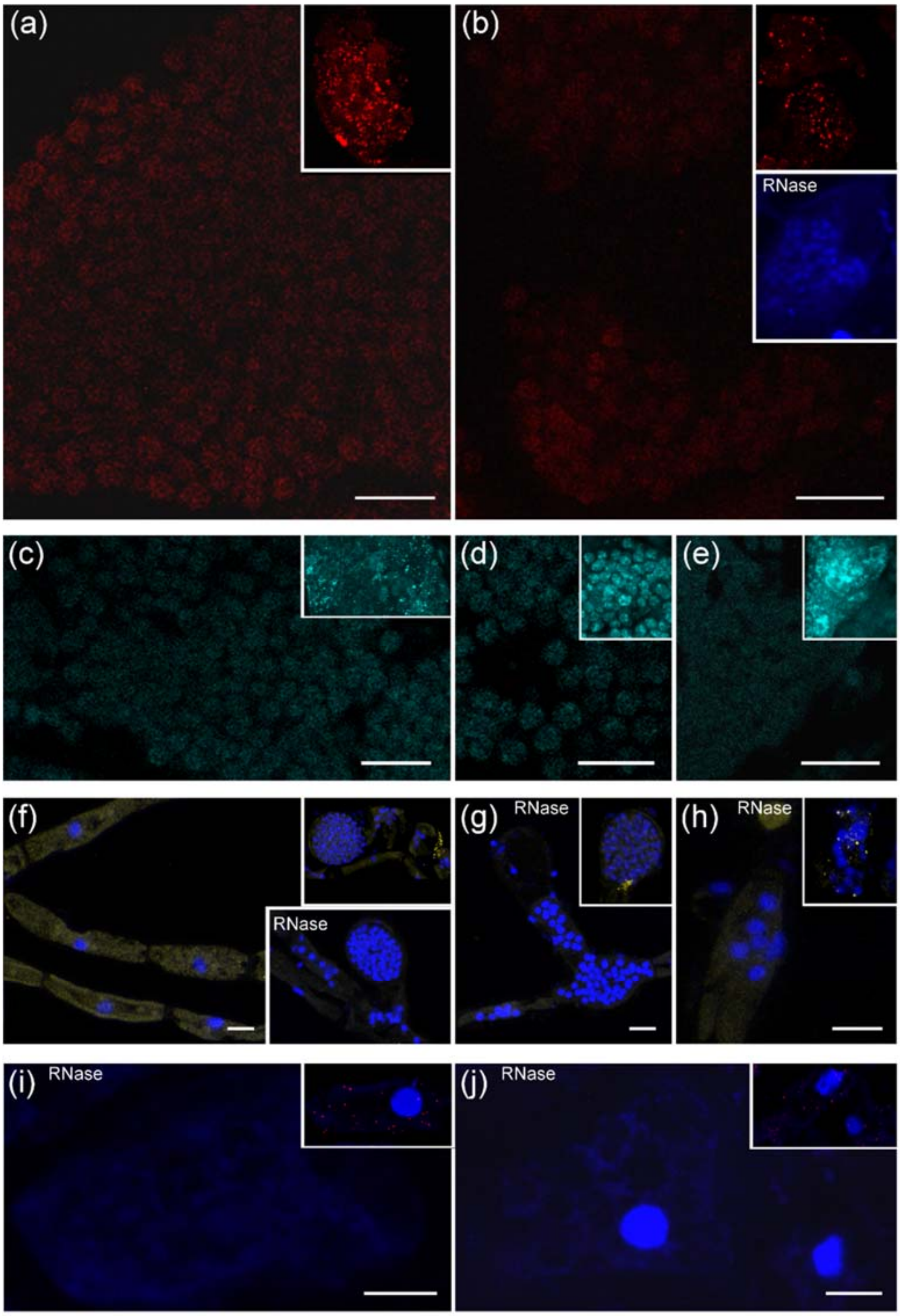


**Figure S5: Negative controls** without padlock/smFISH probe or treated with RNase combined with DAPI staining, unlabeled inserts: corresponding images from Fig. 1. FISH probes should target (a-e) *P. brassicae*, (f-h) *E. siliculosus* (i-j) *Brassica rapa*. a: Negative control of RCA FISH for P. brassicae *PbBSMT* without the padlock probe. b: No probe negative control, smFISH. Insert shows a negative control using the *PbBSMT* probe set after RNase treatment of the tissue. c-e: Negative control of *P. brassicae Actin1* gene detection, RCA FISH without padlock probe. Images show similar development stages as Fig. 1: (c) containing sporulating sporogenic plasmodia, (d) resting spores and (e) in sporogenic plasmodia. f-h: Negative controls for *vBPO* detection experiments, smFISH: (f) uninfected *Ectocarpus* sp. filaments without probes, and (g, h) RNase treatment control of Ec32m infected with *M. ectocarpii*. i, j: RNAse treatment control using brassica *MEX1* probes on infected root cells containing amyloplasts. Bars = 10 µm.

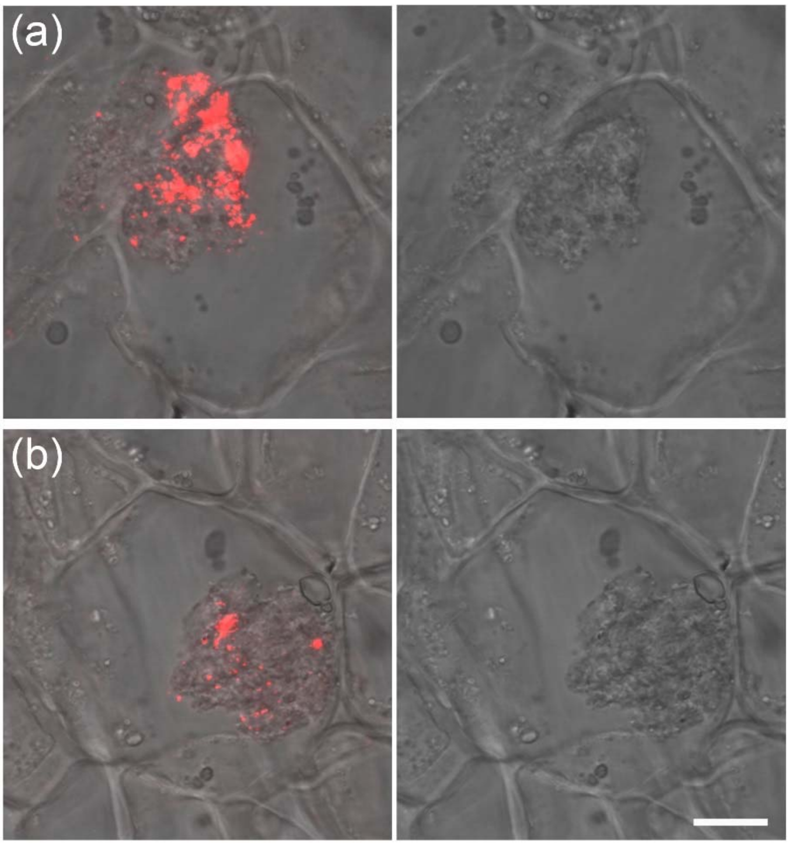


**Figure S6: *PbBSMT* mRNAs can also be localised in young, developing plasmodia.** (a) detection of *PbBSMT* in a young plasmodium appearing to move from one host cell to a not yet infected host cell. Such movement has been described previously, but evidence of cell to cell movement of *P. brassicae* within infected roots is sparse. (b) *PbBSMT* mRNAs in a young plasmodium. These images strengthen the hypothesis, that *PbBSMT* has a masking/protective function against the plant immune system. As this was an isolated observation, these results will need to be confirmed in the future. Images are z-stack maximum projection in young condensed plasmodia (blue is DAPI DNA staining) in Chinese cabbage (gall cross section) host cells using a CLSM. Bars = 10 µm

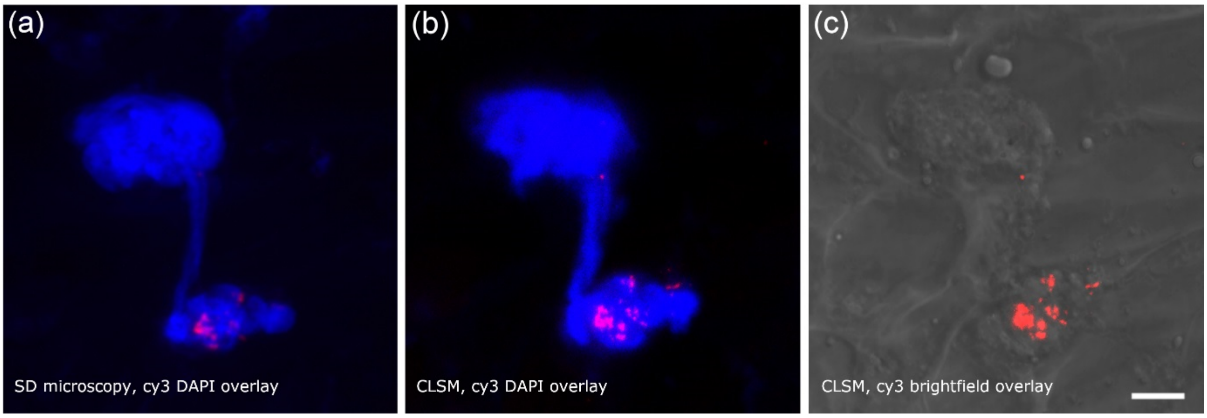


**Figure S7: Comparison of confocal laser scanning microscopy and spinning disk microscopy.** One image was analysed using **(**a) Zeiss Cell Observer.Z1 Spinning Disk (SD) microscope. (b) Leica SP5-II confocal laser scanning microscope (CLSM). (c) shows image (b) with brightfield overlay. The image pattern is identical in both images (a, b). There are clearer and stronger FISH signals detected when using CLSM, but this difference is based on intrinsic methodical differences of CLSM and SD and the camera or detectors used. The image shows a *P. brassicae* sporogenic plasmodium, that appears to move from one cell to the neighbouring cell. The detection of PbBSMT in these mobile plasmodia provides evidence for its role in the suppression of the host defence. Blue = DAPI staining, image was taken from Chinese cabbage clubroots, cross section. Bars = 10 µm

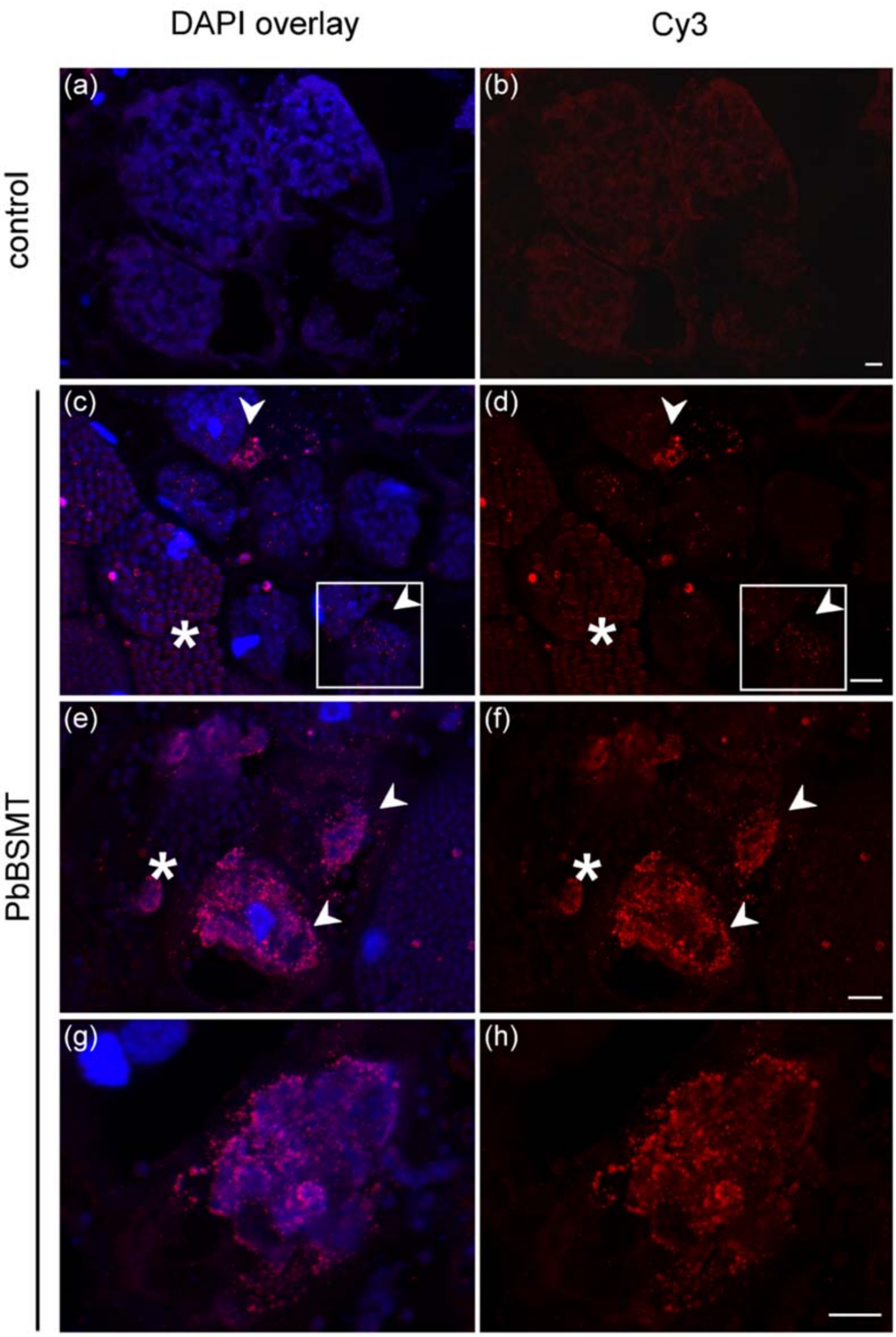


**Figure S8: mRNA localisation of the methyltransferase *PbBSMT* of *P. brassicae* in Chinese cabbage using smFISH.** a, b: control without smFISH probes. Typical signals could not be detected in the lobose, actively growing multinucleate plasmodia and in plasmodia that start to develop into spores. c, d: Overview of multiple P. brassicae infected cells, most of them showing plasmodia in slightly distinct stage during the transition from plasmodial growth to spore formation. The highest number of clear and distinct signals can be seen in cells which are in the beginning of this transition (arrows), while when spore formation has progressed the number and intensity of the signals decreases (asterisks). A selected area is shown in Fig. 2b. e, f: Accumulation of *PbBSMT* mRNAs in plasmodia which start their transformation into resting spores (arrows) and decreased number of mRNAs after spore formation (asterisks). g, h: Another example of *PbBSMT* mRNA accumulation around a plasmodium starting to develop into resting spores. Pictures c-h are presented as maximum projection. Bars =10 µm.

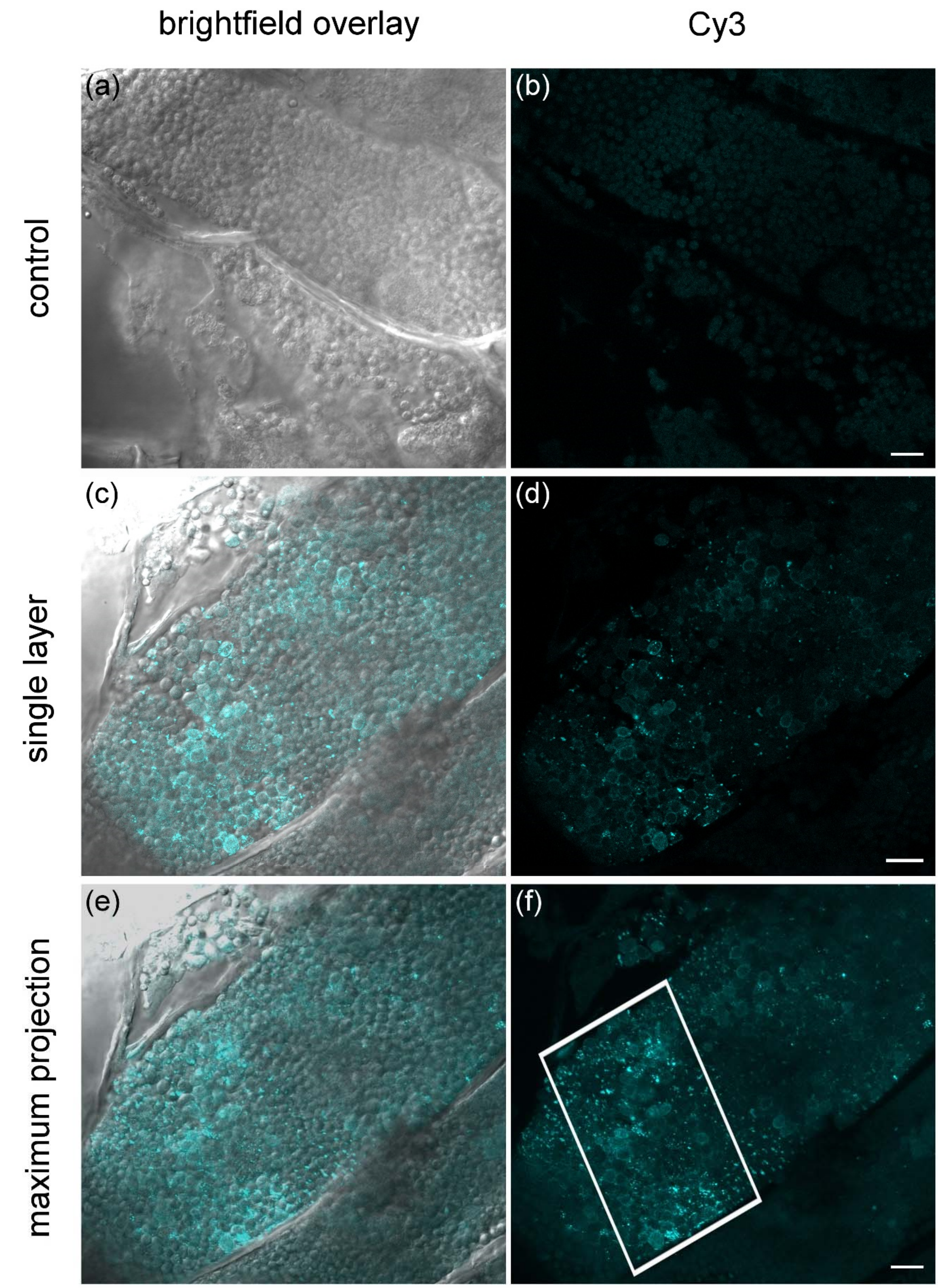


**Figure S9: mRNA localisation of *P. brassicae Actin1* in white cabbage using RCA FISH**. a, b: RCA-FISH control without padlock probe. No signals visible in sporulating sporogenic plasmodia, around and in resting spores. The pictures c-f are an overview of the cell from which a selected area is shown in Fig. 2c: individual z-plane, and e, f: maximum projection of the same cell. d: small, well defined cyan signals distributed across *P. brassicae* which start to get less well defined in panel (f) giving another example of the formation of the apparent signal clustering in z-stack maximum projections and comparison to single z-plane images as discussed in Fig. S3. Bars = 10 μm.

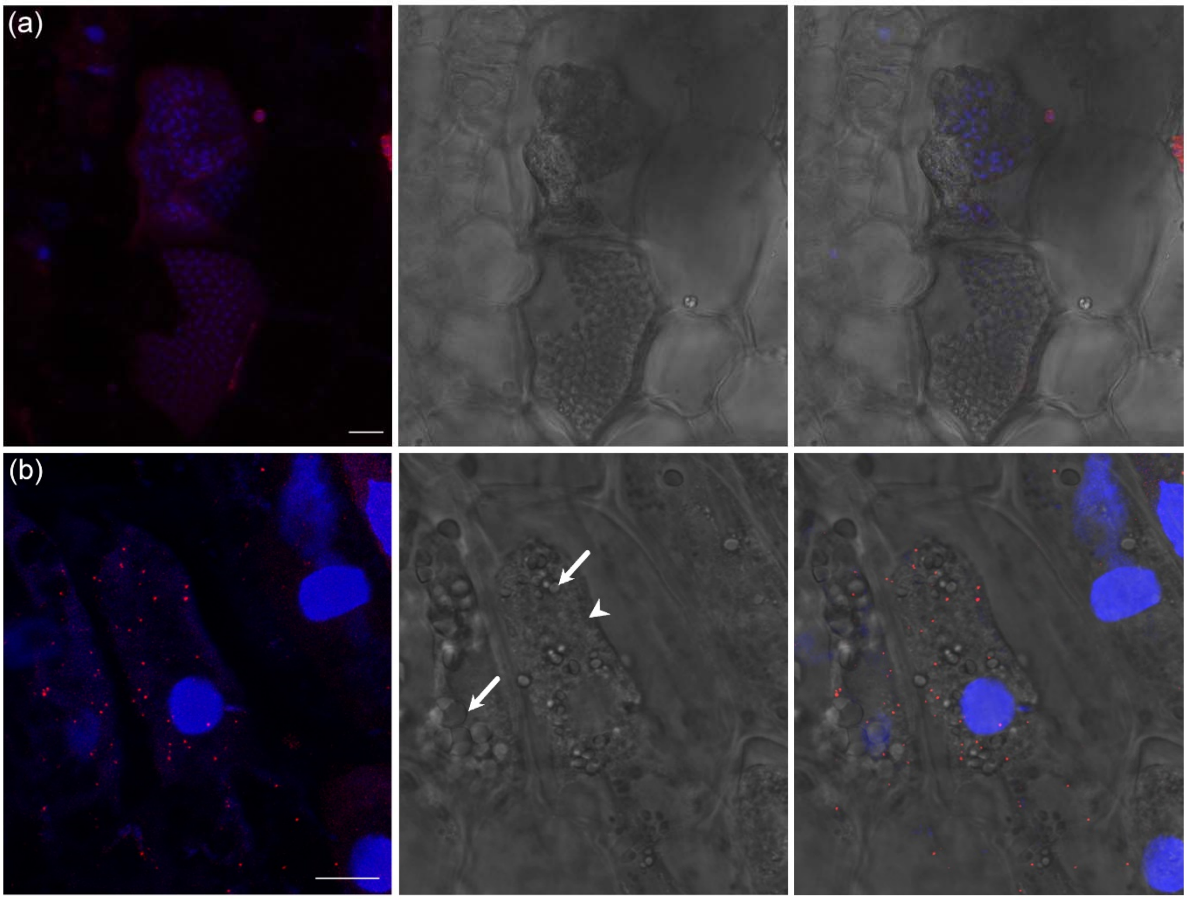


**Figure S10: Detection of the brassica maltose excess protein 1 (*MEX1*) in Chinese cabbage.** a: *MEX1* mRNAs cannot be detected in Chinese cabbage root cells (cross section) without amyloplasts, smFISH. Absence of *MEX1* mRNAs in *Plasmodiophora brassicae* infected host cells (*P. brassicae* plasmodium, arrowhead; resting spores spores, asterisk). b: *MEX1* mRNAs in *Plasmodiophora brassicae* infected host cells (plasmodium, arrowhead), and in not yet infected plant cells containing amyloplasts (arrows). Left panel: overlay pictures with DAPI DNA staining (blue) are shown as maximum projection, right panel: corresponding brightfield images. Bars = 10 µm

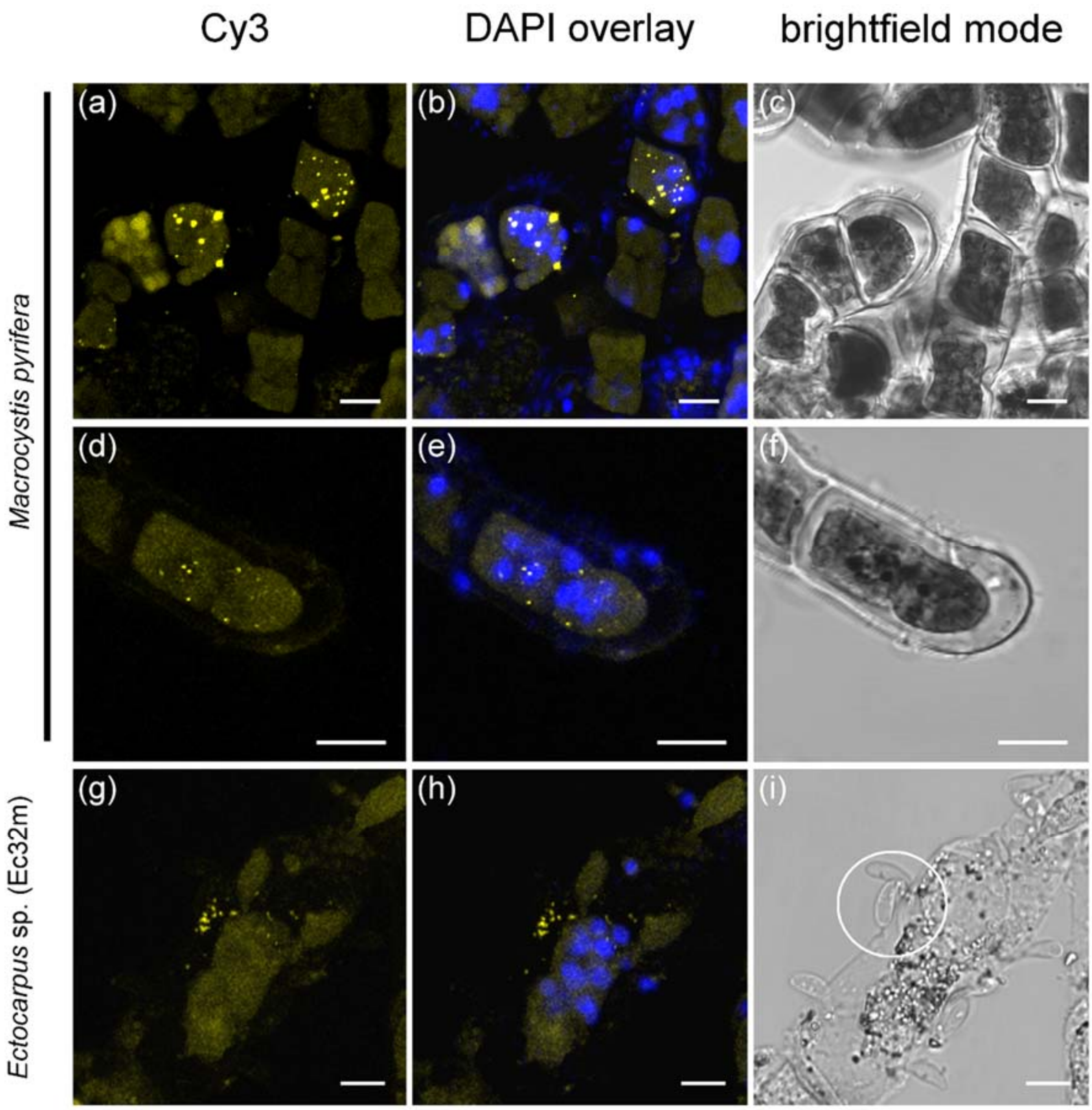


**Figure S11: mRNA localisation of *vBPO* (vanadium-dependent bromoperoxidase) in *Ectocarpus siliculosus* and *Macrocystis pyrifera* infected with *Maullinia ectocarpii*.** Cy3 and DAPI overlay pictures are presented as maximum projection. a-f: yellow *vBPO* signals in infected fresh *M. pyrifera* filaments filled with plasmodia containing several nuclei (DAPI). Signals within filaments and in tips. g-i: yellow *vBPO* signals in infected *Ectocarpus* sp. cells, filled with several pathogen nuclei, where spores (circle) are attached. Bars = 10 µm.

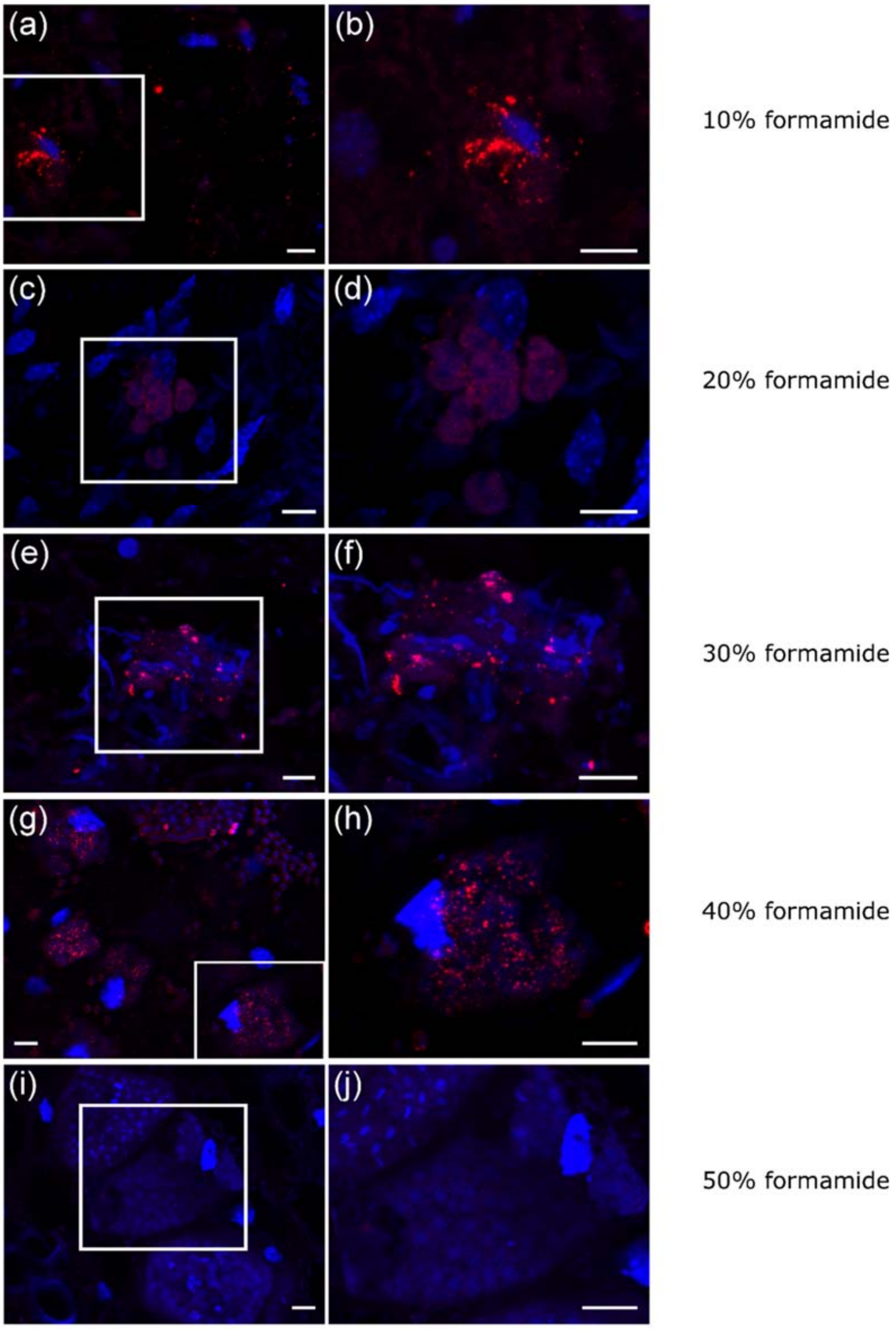


**Figure S12: Testing hybridization efficacy of smFISH probes for *PbBSMT* mRNAs.** a, b: hybridization buffer containing 10% formamide. Positive detection, but mRNA signals are blurred and unclear, b) magnification of the area defined in panel a. c, d: 20% formamide, blurred, unclear signals, worst results in the test series. e, f: 30% formamide, dot-like signals for *PbBSMT*, few signals have the expected size and shape, however many mRNAs still appear to be vast and undefined. g, h: the majority of the detected mRNAs are characterised by clear signals in the expected size and shape. Overall images taken at 40% formamide concentration had the best quality of mRNA detection. i, j: 50% formamide, no mRNAs could be detected . Cross sections of Chinese cabbage root gall infected with *P. brassicae* with mRNA localisation of *PbBSMT* (red signals) with overlay of DAPI channel (blue, DNA). Bars = 10 µm

**Table S1:** smFISH probes for Ec32m *vBPO*. All probes are modified with a Cy3 labelling at the 3′-end.

| **Sequences (5' – 3')** | | | |
|---|---|---|---|
| **1** | gaaggtggcgaagccgac | **20** | gacgtactcgatccgctt |
| **2** | agggaggtgaggatgtgc | **21** | ccgttctgatcgatgcac |
| **3** | cgagtggaggagctggag | **22** | ttcgtacgtgagtccctg |
| **4** | cacttggcgtaccacgcg | **23** | ttggacgccaccttgttg |
| **5** | ttagcacgcggtgcactt | **24** | atgtgagacctgccgatg |
| **6** | cgtgttgtgtaccaggcc | **25** | tccatccggtagtgtacg |
| **7** | gtgtgcagtcgaccggtg | **26** | catcagggcgccgtacac |
| **8** | aggatagtcttgggcagg | **27** | cggacagcactagtctcg |
| **9** | tcagcagctccgtgttct | **28** | ctccgaaaggttgggcag |
| **10** | gttatgcgtcctgactcg | **29** | gaacttgtacgaggccgg |
| **11** | ccgtcgccgttgcatttg | **30** | agaaccttcccgctgtac |
| **12** | atggggagcaggtaggtc | **31** | atgtccttcgcgagcagc |
| **13** | tgaacgggtgacccctcg | **32** | tccgtccgtctagcttga |
| **14** | cgtgtccgcttgggtagg | **33** | tgtgtagagacccttgca |
| **15** | tgtaggcaccaaggttga | **34** | ggtgctcgtcacagaagt |
| **16** | aggaacgccttgagcgtg | **35** | gtgttccagataatggcc |
| **17** | ctctgtccgagctcgaag | **36** | aagtcaccacccttgttc |
| **18** | accaagtccccaaggtag | **37** | tcagagctcgacgtggaa |
| **19** | tgccttcgtcgttggaga | **38** | ccgtcttcattcattgca |

**Table S2:** smFISH probes for *P. brassicae PbBSMT*. All probes are modified with a Cy3 labelling at the 3´-end.

| **Sequences (5' – 3')** | | | |
|---|---|---|---|
| **1** | cggcatcgtggacggata | **25** | gccaatccgactctcttg |
| **2** | gcagcaatggaaacagcc | **26** | aatacgttcccagtcctg |
| **3** | aggctccgggacgtacat | **27** | ggtttcgtctcacggttg |
| **4** | atcacgactccagccatg | **28** | acatagaacatgcgggcc |
| **5** | tagacgcggcgttgtact | **29** | caagatgttgtagcccgg |
| **6** | tcaacgagtgcgcttggc | **30** | gtcatgccaggtgaacgg |
| **7** | aagatctccggcagcatc | **31** | gttagcgttcgttccata |
| **8** | cgtcgactgcgcacatcg | **32** | gttccactactgagaagt |
| **9** | gtagtcggcgatcgtgac | **33** | gaaagactgctcatgcgt |
| **10** | atgtccggccttgtgaac | **34** | gaagtagaccgcctggac |
| **11** | tgctctcgaagatggtca | **35** | cctcgaactcggcattcg |
| **12** | ctttgaccaggacaggcg | **36** | ggtgtcctgcgtagaagg |
| **13** | caggttggtcgatcagga | **37** | acagcgacatcccgtcaa |
| **14** | acatctggcgagtcgtgg | **38** | aacgtccgccaatcgatg |
| **15** | agttgatggctccacagg | **39** | tcatcccaatccgtgttc |
| **16** | cctcgatgtcgtgacgag | **40** | ttccacgcgaacgtgtcg |
| **17** | agacgggttgctttcagc | **41** | gccagaagagaacgacgg |
| **18** | gccgcacaggaagattcg | **42** | cgtctgccagatattgct |
| **19** | ggaagcacctgctcgaag | **43** | gcaatgtggtgacgtgct |
| **20** | ggcgatgtcgagaattcc | **44** | gatatcgtcgagcaggcg |
| **21** | cattgcagagccgatccg | **45** | ttgtacaccagcgggttg |
| **22** | atatggggacgctgcgat | **46** | gtatgcctgaagccagtt |
| **23** | cccagaaagccgatttca | **47** | tcagtgtttcgcgatggt |
| **24** | agcgaccgaaacgtcgtg | **48** | gacagtgtaccaatacca |

**Table S3:** smFISH probes for *B. rapa MEX1*. All probes are modified with a Quasar570 labelling at 3′ -end.

| **Sequences (5' – 3')** | | | |
|---|---|---|---|
| **1** | gacattgcgatggttttagc | **25** | aaaatcttcccacaatcgcc |
| **2** | atcaacgagcggttactagc | **26** | aacacagagtccaccaacag |
| **3** | ggaagagcgaggagaagtcg | **27** | tagaccacatgacttgaggt |
| **4** | ggcgggagaaacgagatgta | **28** | ctgtttggtacaaaagggac |
| **5** | tatacaagcacgcttcagcg | **29** | aacaaaagcagttgtcccag |
| **6** | aatcggattccaccgagtca | **30** | taaatacagccactacagca |
| **7** | cgagtcgtcggcgattcgaa | **31** | tgggagtttctcggttcgag |
| **8** | gagttagttgcacgaacagg | **32** | ggacaaacttaacaccttcc |
| **9** | tctctaaacctagagatcct | **33** | gctgtccatccagataaaga |
| **10** | ccattcttcgtattccttaa | **34** | gcatccacatgaacataagg |
| **11** | gagaatttcgcagtccatga | **35** | tttgtccacatttgggaaac |
| **12** | aaacggaatattcgctccgc | **36** | tatgttgtctggacttagaa |
| **13** | tgatctgcggcaattgaagc | **37** | ttgtgattggtgataagcct |
| **14** | cgccaaaagattctgtgcat | **38** | ttccccgtcatcgaaagcaa |
| **15** | atggaacagccgaaagtgcg | **39** | ataatgctcttgggagcata |
| **16** | aaaccagtcaacatccccag | **40** | gaaccacatcaaatcacgga |
| **17** | aagcaacgaaaggtttccta | **41** | gagagttgcccatattgaac |
| **18** | tttctctcttctttgcgaaa | **42** | cacagaatgtttccatatcc |
| **19** | gtttgcacaatagctgcttc | **43** | ctcgtgcagttagacatgta |
| **20** | gtgagtggagatgactccaa | **44** | ttgtggatgctgcgaagaat |
| **21** | atggtgagctgcgaaaggac | **45** | cctatccatgagatcaaacc |
| **22** | aaacggcaaaggcatagctc | **46** | gctgaatctctccataaagc |
| **23** | caacagccgaagtagcaaca | **47** | aacggagagttgtgaccatg |
| **24** | gtcgtactgagcttaccaaa | **48** | acccaaaaaccaactccttc |

**Table S4:** z-stack thickness, number of slices, and z-stepsize used in the images of this work. All images were recorded with minimum pinhole settings, resulting in a slice thickness of 0.85µm with the 63x objective.

| Figure 2 | Thickness (µm) | planes | slice thickness (µm) |
|---|---|---|---|
| a | 32,60 | 38 | 0,86 |
| b | 25,51 | 33 | 0,77 |
| c | 19,13 | 39 | 0,49 |
| d | 14,98 | 22 | 0,68 |
| e | 16,74 | 22 | 0,76 |
| f | 21,15 | 29 | 0,73 |
| g | 9,41 | 17 | 0,55 |
| h | 14,27 | 21 | 0,68 |
| i | 19,13 | 39 | 0,49 |
| j | 20,73 | 39 | 0,53 |
| Figure 3 | | | |
| d, g | 17,72 | 31 | 0,57 |
| e, h | 20,73 | 27 | 0,77 |
| f, i | 9,27 | 14 | 0,66 |
| Figure S1 | | | |
| a | 17,72 | 31 | 0,57 |
| b | 20,73 | 27 | 0,77 |
| c | 32,60 | 38 | 0,86 |
| d | 25,51 | 33 | 0,77 |
| e | 29,50 | 38 | 0,78 |
| Figure S2 | | | |
| a | 35,25 | 41 | 0,86 |
| b | 15,95 | 21 | 0,76 |
| c | 34,28 | 44 | 0,78 |
| d | 25,51 | 33 | 0,77 |
| e | 26,31 | 34 | 0,77 |
| f | 16,74 | 20 | 0,84 |
| g | 22,03 | 26 | 0,85 |
| h | 12,76 | 17 | 0,75 |
| i | 9,27 | 14 | 0,66 |
| Figure S3 | | | |
| | 32,60 | 38 | 0,86 |
| Figure S4 | | | |
| | 15,15 | 20 | 0,76 |
| Figure S5 | | | |
| b | 42,3 | 43 | 0,98 |
| b RNase | 21,15 | 43 | 0,49 |
| f | 8,22 | 15 | 0,55 |
| f RNase | 14,6 | 13 | 1,12 |
| g | 17,62 | 36 | 0,49 |
| h | 18,13 | 37 | 0,49 |
| i | 26,39 | 38 | 0,69 |
| j | 28,53 | 41 | 0,70 |
| Figure S6 | | | |
| a | 12,92 | 45 | 0,29 |

| | | | |
|---|---|---|---|
| b | 11,46 | 40 | 0,29 |
| Figure S7 | | | |
| | 10,28 | 36 | 0,29 |
| Figure S8 | | | |
| a, b | 42,3 | 43 | 0,98 |
| c, d | 25,51 | 33 | 0,77 |
| e, f | 22,03 | 26 | 0,85 |
| g, h | 16,74 | 20 | 0,84 |
| Figure S9 | | | |
| | 19,13 | 39 | 0,49 |
| Figure S10 | | | |
| a | 24,55 | 40 | 0,61 |
| b | 19,13 | 39 | 0,49 |
| Figure S11 | | | |
| a, b, c | 15,15 | 20 | 0,76 |
| d, e, f | 14,27 | 21 | 0,68 |
| g, h, i | 7,64 | 14 | 0,55 |
| Figure S12 | | | |
| a | 27,19 | 55 | 0,49 |
| b | 15,95 | 39 | 0,41 |
| c | 11,75 | 29 | 0,41 |
| d, e | 26,31 | 34 | 0,77 |
| f | 19,97 | 35 | 0,57 |
