## Supplementary material for "Biotrophic interactions disentangled: *In situ* localisation of mRNAs to decipher plant and algal pathogen – host interactions at the single cell level": Supplemtn image analysis

**Additional information for raw data and image processing:**

Numbers of the folders correspond to the images included in the text. In the manuscript and the supplementary material all figures are shown as z-stack maximum projections which were generated from the raw image files using ImageJ (version 2.0.0; Java 1.8.0). Linear brightness and contrast adjustments on these maximum projections were made in GIMP 2.8.22 (www.gimp.org). All images were taken in three subsequently recorded excitation and emission settings (Fig.1) regardless of the used fluorophores. Where the biological material permitted, brightfield images were generated in addition to the fluorescent images.

In the raw data files images with the settings excitation: 514 nm and emission: 550 - 585 nm are labelled as “red” channel (because the emission corresponds to the red colour spectrum), images with the settings excitation: 488 nm and emission: 500 - 550 nm are labelled as “green” channel (because the emission corresponds to the green part of the colour spectrum), images with the settings excitation: 405 nm and emission: 430 - 485 nm are labelled as “blue” channel (because the emission corresponds to the blue part of the colour spectrum). Plant and algal material often has a lot of autofluorescence itself, therefore we aimed to use settings that did yield in clear mRNA detection signals while reducing the autofluorescent background. On the first sight the images can, therefore, appear to be underexposed. As illustrated in the images below, this was done to make as clear images as possible from the beginning to reduce the amount of image manipulation. Also the negative controls, treatment without FISH probes or RNase treatment, were analysed with the same settings. They were adjusted for the minimum background fluorescence to reduce potential artefact background signals.

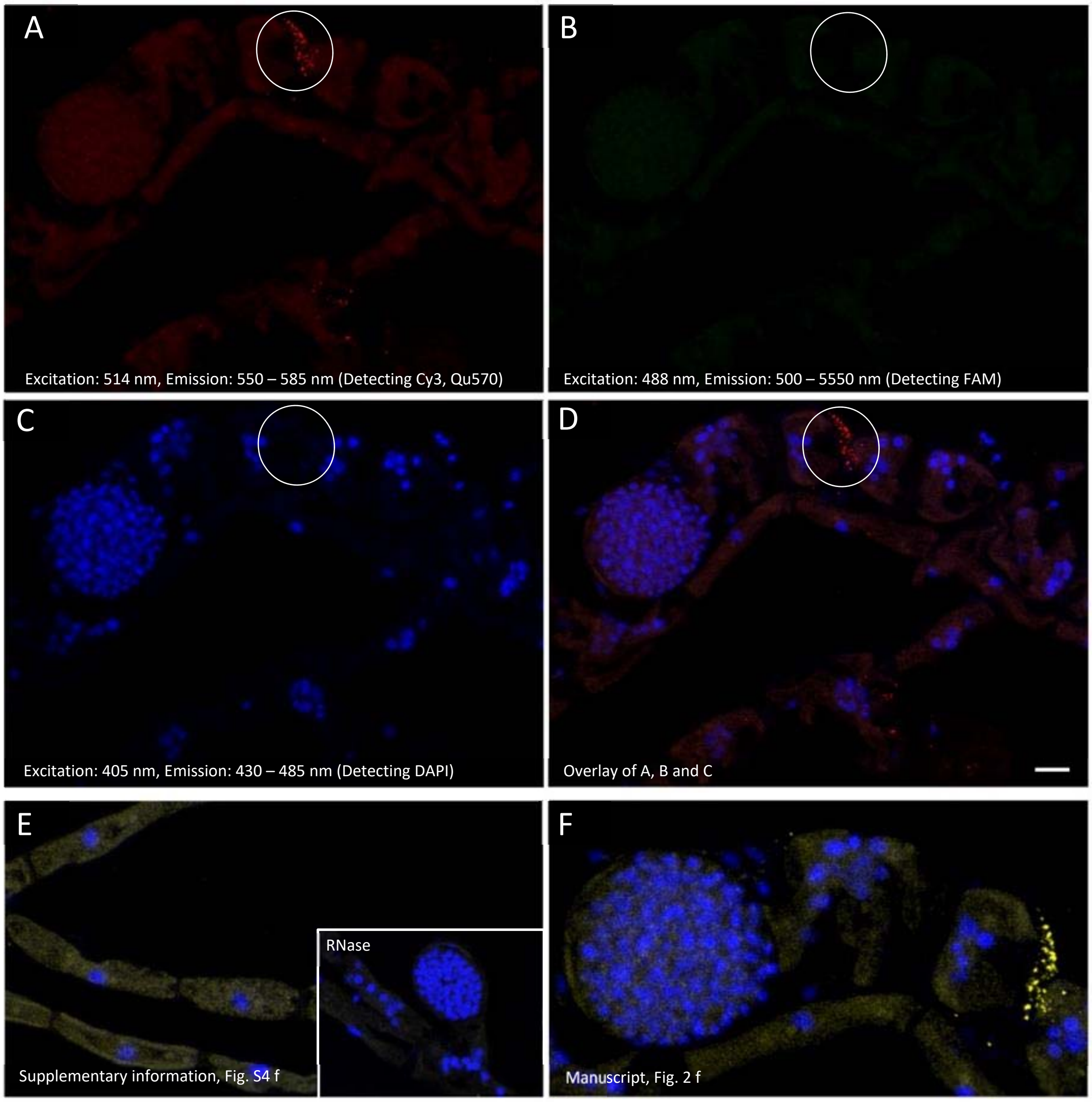


**Fig. 1: z-stack maximum projection without any brightness/contrast adjustments. *M. ectocarpii* infected *E. siliculosus* filaments with smFISH *vBPO* mRNA probe (visible in 1A, red dots highlighted by the circle).** A-C: sample with Cy3 probe (red dots, circle) analysed in the red channel (A), green channel (B) and blue channel (C). D: Overlay of all 3 channels. E: negative controls, without probe or after RNase treatment, as presented in the Supplementary Information (Fig. S4). F: Image as presented in the main manuscript (Fig. 2 f). Scale bar = 10 µm

Because we chose to use identical recording settings for all images (i.e. image the same excitation/emission settings for each experiment regardless of the probe), sometimes the mRNA signals can have a slight carry over between the different settings. We tried to narrow the range of the measured wavelength (Fig. 2) for best results, minimizing the shine-through, but it was impossible to exclude it completely. However, we did select those settings regardless of this slight signal carry over, because our samples did have the strongest autofluorescence when those excitation and emission spectra were recorded, which helped us greatly to identify and remove artefacts from our analysis.

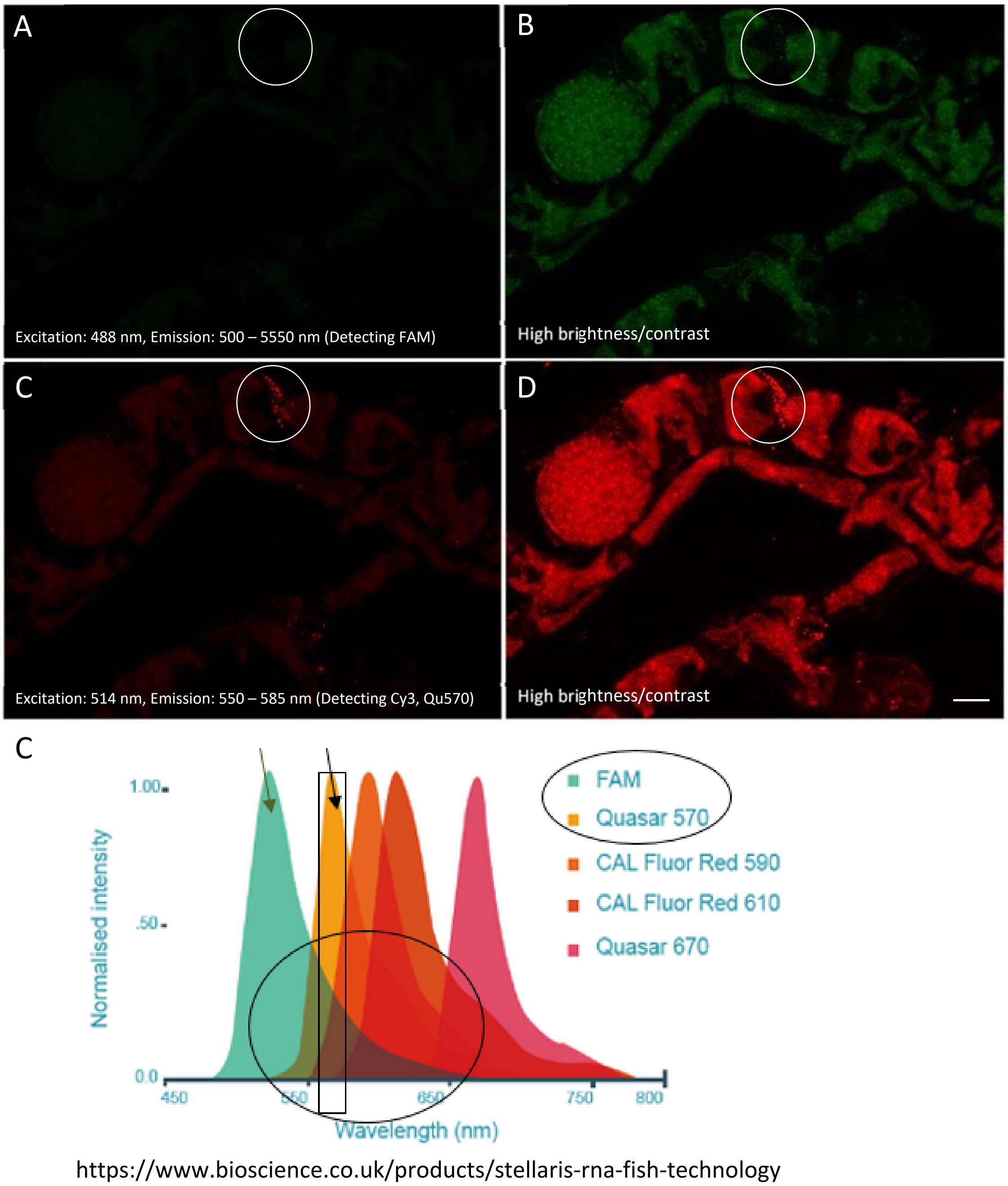


**Fig. 2: Light "shine-through signals" appear in the green channel after brightness/contrast adjustment, although there is no FAM probe due to spectra overlay.** A, C: Raw data image shown as z-stack maximum projection without any brightness/contrast adjustments. B, D: Same image, with highly increased brightness/contrast adjustments. (Brightness: 110, contrast: 105; GIMP) Light "shine-through signals" are slightly visible (circle) in the green channel, without FAM probe. C: different spectra of different dyes, showing the overlay (circle) of FAM (green arrow) and Quasar570 (Cy3) (black arrow) spectra. Scale bar = 10 µm.

In selected cases, image deconvolution was performed. Raw image stacks were deconvoluted with the Huygens software package (Scientific Volume Imaging, The Netherlands). The differences between deconvolution and raw data images of DAPI-Cy3 overlay images is shown in Fig. 3.

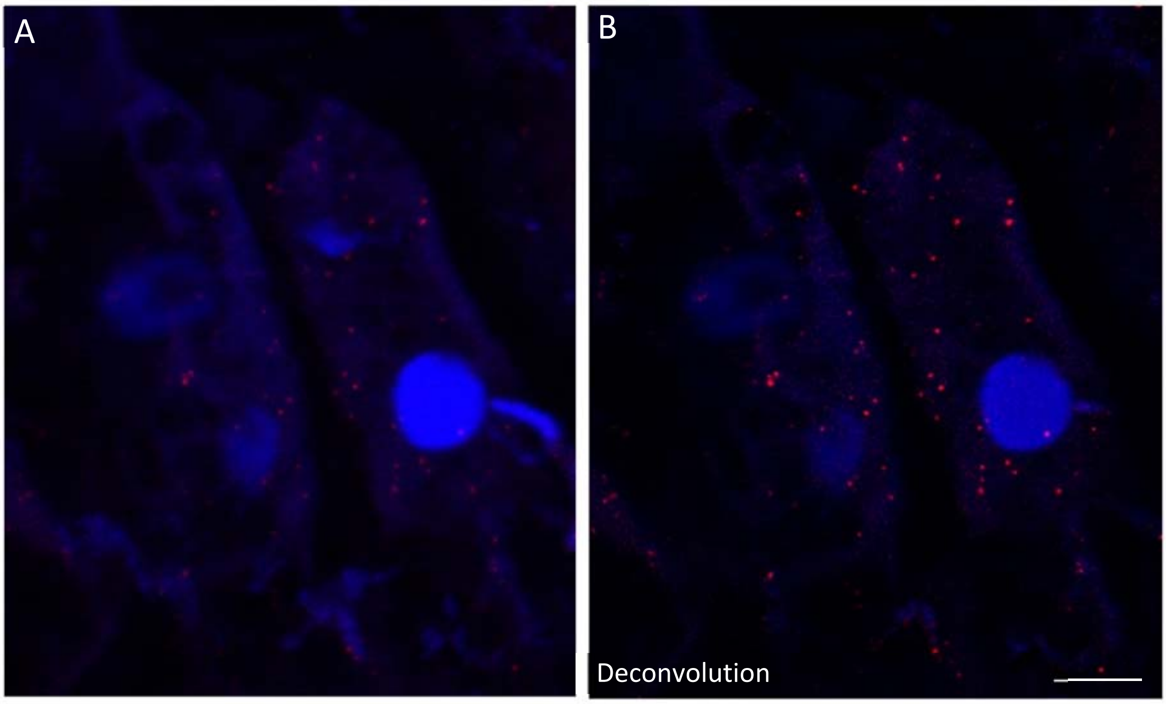


**Fig. 3: Comparison between deconvolution and no deconvolution of raw data images.** It shows *MEX1* mRNAs in the cytosol of *P. brassicae* infected *B. rapa* root cells using smFISH (red signals, Quasar 570, blue: DAPI DNA staining). A: Overlay of DAPI and Cy3 channel. Without deconvolution. B: Overlay of DAPI and Cy3 channel with deconvolution. Scale bar = 10 µm.
